# Notch signaling represses cone photoreceptor formation through the regulation of retinal progenitor cell states

**DOI:** 10.1101/2020.11.09.375832

**Authors:** Xueqing Chen, Mark M. Emerson

**Affiliations:** Biology PhD Program, The Graduate Center, The City University of New York, New York, NY, 10016; Department of Biology, The City College of New York, The City University of New York, New York, NY, 10031; Biochemistry PhD Program, The Graduate Center, The City University of New York, New York, NY, 10016

**Keywords:** retinal progenitor cells, multipotency, cis-regulation, cone photoreceptors, Notch inhibition, dominant-negative, chicken, electroporation

## Abstract

Notch signaling is required to repress the formation of vertebrate cone photoreceptors and to maintain the proliferative potential of multipotent retinal progenitor cells. However, the mechanism by which Notch signaling controls these processes is unknown. Recently, restricted retinal progenitor cells with limited proliferation capacity and that preferentially generate cone photoreceptors have been identified. Thus, there are several potential steps during cone genesis that Notch signaling could act. Here we use cell type specific cis-regulatory elements to localize the primary role of Notch signaling in cone genesis to the formation of restricted retinal progenitor cells from multipotent retinal progenitor cells. Localized inhibition of Notch signaling in restricted progenitor cells does not alter the number of cones derived from these cells. Cell cycle promotion is not a primary effect of Notch signaling but an indirect effect on progenitor cell state transitions that leads to depletion of the multipotent progenitor cell population. Taken together, this suggests that the role of Notch signaling in cone photoreceptor formation and proliferation are both mediated by a localized function of Notch in multipotent retinal progenitor cells to repress the formation of restricted progenitor cells.

## Introduction

Notch signaling is an evolutionarily conserved signaling pathway that functions in diverse cellular contexts to drive both cell cycle and cell fate decisions. In canonical Notch signaling, extracellular binding of a member of the Delta/Serrate/Lag family of proteins to the extracellular portion of the transmembrane Notch receptor leads to a proteolytic cleavage that releases the Notch intracellular domain (NICD) from the membrane (Bray, 2006; Kopan and Ilagan, 2009). The NICD then translocates to the nucleus, and through a primary interaction with the MAML protein, forms a tripartite complex with the Rbpj transcription factor that affects transcription of target genes (Mumm and Kopan, 2000; Wu et al., 2000). Notch signaling can act iteratively and participate in several distinct decisions that occur during the process of formation of a particular cell type. For example, during the formation of Drosophila sensory neurons from ectoderm, Notch signaling is critical in three successive cell fate decisions (Artavanis-Tsakonas et al., 1991; Bray, 1998).

During the formation of the vertebrate retina, a functional role for Notch signaling in both proliferation and in cell fate choice has been identified (Hayes et al., 2007; Henrique et al., 1997; Jadhav et al., 2006a; Mizeracka et al., 2013a; Nelson et al., 2007). Induction of conditional loss-of-function alleles for the Notch1 receptor during early retinogenesis leads to cell cycle exit and increased cone photoreceptor formation (Jadhav et al., 2006b; Yaron et al., 2006). In one of these studies, a loss of horizontal cells (HCs) was also described (Yaron et al., 2006). Similarly, conditional Rbpj loss-of-function increases the formation of cone photoreceptors while also promoting the formation of retinal ganglion cells (RGCs) (Riesenberg et al., 2009). Both Notch1 and Rbpj loss-of-function conditions result in decreased proliferation. Conditional loss of Notch1 in the later postnatal period, after the window of cone genesis, reveals that rod photoreceptors are also increased, which suggests that Notch signaling plays a more general role in repression of the photoreceptor fate (Jadhav et al., 2006b). In addition, it has been suggested in this postnatal context that Notch signaling can function in postmitotic cell fates to repress the rod fate (Mizeracka et al., 2013b).

While a role for Notch signaling in the generation of cone photoreceptors has been identified, the pan-retinal inhibition of Notch signaling through chemical inhibition and conditional alleles has not allowed for a specific role of Notch to be determined. Recently, the formation of cones has been identified as a multistep process. During the early stages of retinal formation when cone photoreceptors are preferentially generated, multipotent retinal progenitor cells (RPCs) divide to form restricted (also known as neurogenic RPCs) RPCs that can be identified by the activity of Thrb cis-regulatory elements, the expression of specific transcription factors (Otx2, OC1 and Olig2), and the decreased expression of canonical RPC markers, such as Vsx2 (Buenaventura et al., 2018; Emerson and Cepko, 2011; Emerson et al., 2013; Hafler et al., 2012; Jean-Charles et al., 2018; Suzuki et al., 2013). These restricted RPCs lack the diverse cell fate potential of multipotent RPCs, and instead, preferentially divide to form cone and HCs as well as a minor number of RGCs (Cepko, 2014; Emerson et al., 2013; Schick et al., 2019). Thus, Notch signaling could mediate cone genesis through regulation of the multipotent to restricted RPC transition, the cell fate choices of the restricted RPC daughter cells, or in the production of cones directly from multipotent RPCs or other cell types.

In this study, we used defined cis-regulatory elements to both quantitate the effects of Notch signaling inhibition and also to limit interference with Notch signaling to specific cell types. We determined that during the formation of cone photoreceptors, inhibition of Notch signaling specifically promoted the formation of restricted RPCs from multipotent RPCs but did not promote the cone fate in the daughter cells of restricted RPCs. As expected for such a shift in cycling populations, there was minimal short-term effect on proliferation as measured by EdU labeling. Taken together, this suggests that a primary role of Notch signaling during cone photoreceptor genesis is to regulate the formation of restricted RPC states from multipotent RPCs.

## Results

### Evaluation of Notch reporter activity in response to MAML dominant-negative

To provide cell-type specific precision and rapid interference with Notch signaling, we used a dominant-negative approach. We first created a chicken version of a previously validated, and widely used, C-terminal truncation version of MAML that retains the ability to interact with the Notch intracellular domain but lacks the domains necessary to induce a transcriptional response (referred to as MAML-DN) (Tan et al., 2016; Weng et al., 2003). The chicken MAML1 gene was used, as it was the MAML gene most highly expressed in the developing chicken retina according to reported RNA-Seq data (Buenaventura et al., 2018). The reported MAML-DN construct encoded a fusion protein with the N-terminal region of MAML placed N-terminal to GFP. However, we expected to use GFP reporters as an output in our experiments, and so engineered a fusion protein of the MAML N-terminal region with GAPDH instead of GFP. Two AU1 tags were added to aid in tracking the expression of the protein (Fig 1A) (Shevtsova et al., 2006).

**Fig. 1.**
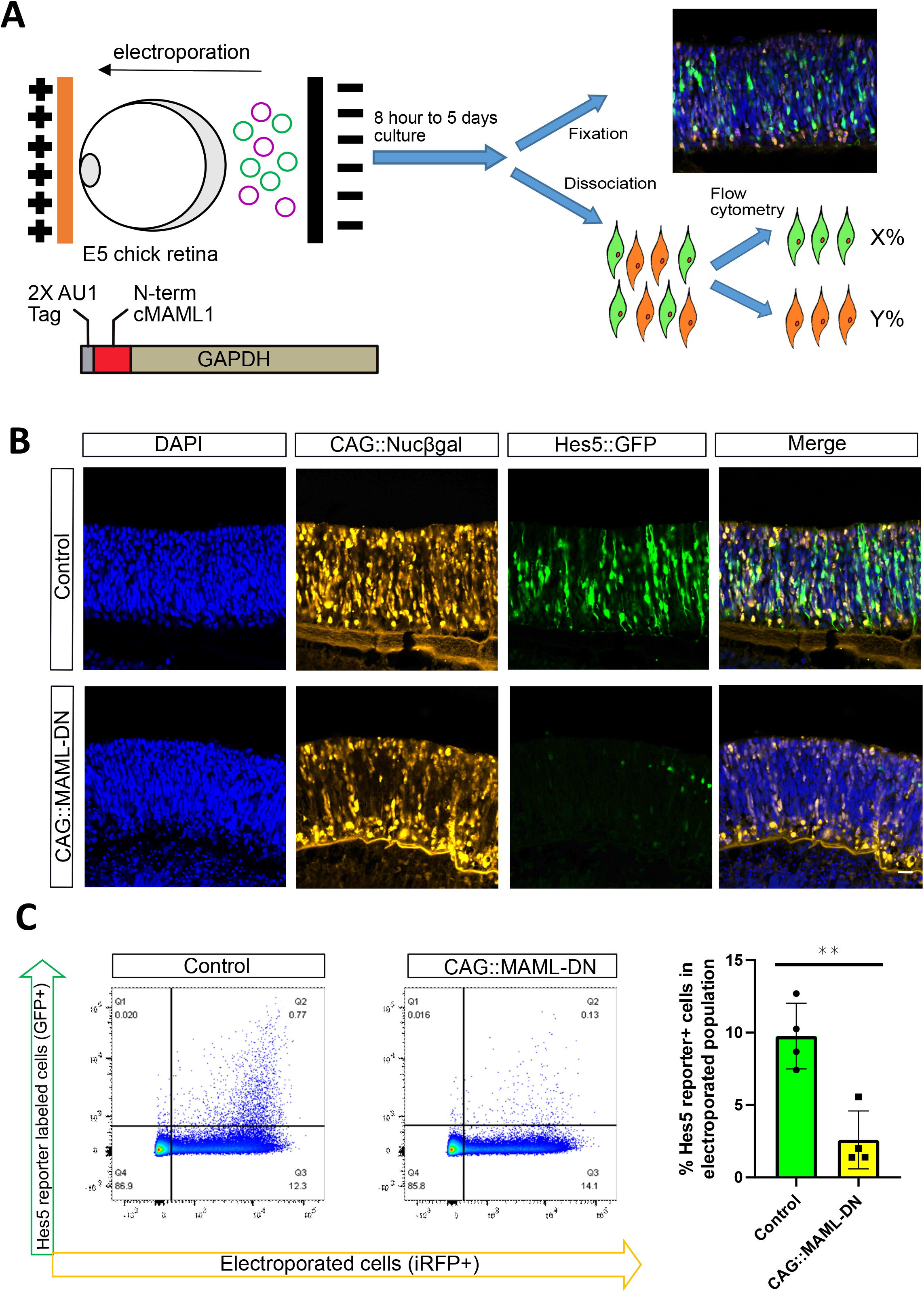
CAG::MAML-DN inhibits the expression of the Hes5 Notch reporter **A**. Schematic representation of the electroporation paradigm with plasmids introduced ex vivo into E5 chick retinas, cultured for two days, dissociated into single cells and analyzed by flow cytometry. A schematic of the MAML dominant-negative construct is shown. **B**. Confocal images of vertically sectioned E5 chick retinas co-electroporated with CAG::Nucßgal, Hes5::GFP Notch reporter, and with or without CAG::MAML-DN and then cultured for two days. The scale bar shown in the bottom right picture denotes 40 µm and applies to all images. All images are oriented with the scleral side of the retina at the top of the image. **C**. Flow cytometry dot plots of dissociated chick retinal cells electroporated *ex vivo* at E5 with a co-electroporation control (CAG::iRFP), Hes5::GFP, and with or without CAG::MAML-DN. The bar graph on the right shows the percentage of Hes5::GFP reporter-positive cells within the electroporated population. The Shapiro-Wilk normality test was used to confirm the normal distribution. ** signifies p<0.01 with a two-tailed student’s t-test. Each point represents one biological replicate.

To test the effectiveness of the MAML-DN construct, we used a previously validated chicken Notch reporter - Hes5::GFP (Vilas-Boas et al., 2011). This reporter uses both transcriptional regulatory elements and 3’ untranslated regions of the chicken Hes5-1 gene, which was previously identified as a target of Notch signaling during chicken development (Fior and Henrique, 2005).

E5 chicken retinas were co-electroporated ex vivo with Hes5::GFP, a ubiquitous electroporation control (CAG::Nucßgal), and with or without the MAML-DN construct. After two days of culture, retinas were imaged by confocal microscopy or analyzed by flow cytometry (Fig. 1A). In control retinas, the Hes5::GFP reporter was active primarily in the neuroblast layer of the retina, where multipotent RPCs are located, and the reporter activity was drastically abrogated in response to MAML-DN introduction (Fig. 1B). Strikingly, we found in all the experimental groups that the retinal distribution of the electroporated population was altered, with cells found predominantly in the scleral region where photoreceptor cells and Otx2/OC1+ RPCs are enriched, as well as the RGC layer, but not in the middle portion of the retina where most multipotent RPCs are localized (Fig. 1B) (Buenaventura et al., 2018). To test the specificity of the effects observed with this MAML-DN construct, a plasmid encoding an AU1-GAPDH protein without the MAML N-terminal region was used in the same experimental workflow. A typical distribution of cells and levels of the Hes5::GFP reporter were observed with this control construct (Fig. S1A). Immunofluorescent detection of the AU1 tag revealed a predominantly cytoplasmic distribution for the AU1-GAPDH construct that appeared more nuclear in the MAML-DN version (Fig. S1A).

To quantify Notch reporter activity, we used a flow cytometry assay and calculated the percentage of GFP-positive cells within the electroporated cell population (Fig. 1C). The quantification supported the conclusions of the confocal imaging results, as Hes5::GFP reporter activity showed a significant decrease in response to MAML-DN introduction. The negative control AU1-GAPDH vector did not interfere with the Hes5::GFP activity (Fig. S1B). To further confirm this effect, we employed an alternative dominant-negative strategy in which we N-terminally or C-terminally fused the repressor domain of engrailed to the full-length Rbpj protein coding sequence (EnR-Rbpj or Rbpj-EnR). This modification would be predicted to convert Rbpj into a dedicated repressor that would continually repress Notch target genes, even under conditions of active Notch signaling (Vickers and Sharrocks, 2002). In addition, we created a plasmid that would express a Rbpj protein with a R218H mutation, which was previously characterized as a dominant-negative form of Rbpj (Chung et al., 1994). Hes5 reporter activity was significantly decreased in the presence of all four dominant-negative constructs, thus identifying multiple dominant-negative approaches to inhibit Notch signaling (Fig. 1 and Fig. S2).

### Inhibition of Notch promotes the formation of restricted progenitor cells

A mouse knockout of the Notch1 receptor leads to an increase in cone photoreceptors (Jadhav et al., 2006b; Yaron et al., 2006), but the mechanism behind this phenotype is unknown. One possibility is that in the knockout, multipotent RPCs produce more of the RPC type that is restricted to cone and HC fates, which in turn divide to form cones. The cis-regulatory element ThrbCRM1 provides a robust tool to answer the question as it is active specifically in this restricted RPC type (Emerson et al., 2013).

We first tested the effect of Notch inhibition on specific cell types using DNA reporters, including reporters with enrichment in multipotent RPCs (VSX2ECR4; Buenaventura et al., 2018), cone/HC restricted RPCs (ThrbCRM1; Emerson et al., 2013), subsets of cones (ThrbCRM2; Emerson et al., 2013; Schick et al., 2019) and HCs/RGCs (OC1ECR22; Patoori et al., 2020; Schick et al., 2020). E5 chicken retinas were co-electroporated with an electroporation control, with or without the MAML-DN, and assessed by confocal microscopy after two days culture. In retinas in which MAML-DN was introduced, the number of ThrbCRM1 reporter-labeled restricted RPCs was visibly increased, while the number of VSX2ECR4-labeled multipotent RPCs was decreased and presented with altered morphology (Fig. 2A). The number of ThrbCRM2 reporter-labeled cones and OC1ECR22 reporter-labeled HCs/RGCs were qualitatively unchanged at this timepoint (Fig. S3A). The quantification of these cell types using flow cytometry statistically confirmed these effects (Fig. 2B). In addition, in retinas in which VSX2ECR4::GFP and ThrbCRM1::TdTomato were co-electroporated, the number of GFP/TdTomato double-positive cells dramatically increased from the low amount that is typically observed (Buenaventura et al., 2018). Taken together, this suggests that the broad inhibition of Notch signaling leads to the formation of supernumerary ThrbCRM1 cells and these new cells are derived from multipotent RPCs. To confirm the specificity of the MAML-DN effect on ThrbCRM1, we used the AU1-GAPDH as a negative control and observed no significant change in ThrbCRM1::GFP levels (Fig. S1B). Intriguingly, ThrbCRM2-labeled cones did not significantly increase after 2 days culture, which could indicate a delay in cone genesis at this time point.

**Fig. 2.**
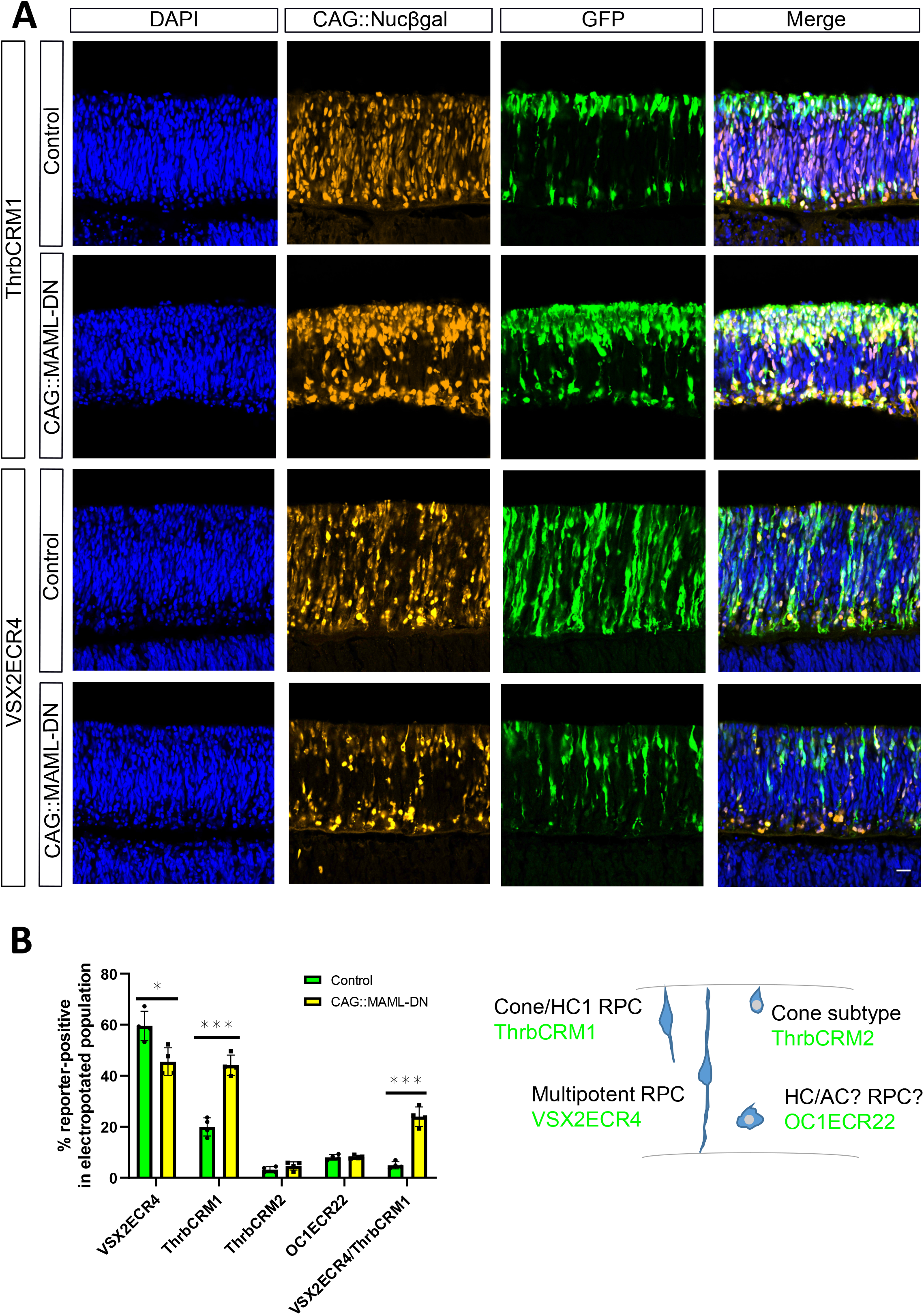
CAG::MAML-DN induced Notch inhibition promotes ThrbCRM1 restricted RPC formation **A**. Confocal images of vertically sectioned E5 chick retinas co-electroporated with CAG::Nucßgal, either ThrbCRM1 or VSX2ECR4 GFP reporters, and with or without CAG::MAML-DN and then cultured for two days. The scale bar shown in the bottom right picture denotes 40 µm and applies to all images. All images are oriented with the scleral side of the retina at the top of the image. **B**. Flow cytometry quantification of the percentage of ThrbCRM1::TdTomato, ThrbCRM2::TdTomato, VSX2ECR4::GFP and OC1ECR22::GFP reporter-positive cells within the CAG::iRFP electroporated population in E5 retinas cultured for two days. Green bars represent control conditions and yellow bars inclusion of the CAG::MAML-DN construct. The Shapiro-Wilk normality test was used to confirm the normal distribution. * signifies p<0.05, ** signifies p<0.01, *** signifies p<0.001 with a two-tailed student’s t-test. Each point represents one biological replicate.

The increase in ThrbCRM1 reporter-positive cells could be due to an increase in the ThrbCRM1 RPC state, but it could also represent an increase in their daughter cell production (cones, HCs, or RGCs) that could be missed by the specific reporters used (Schick et al., 2019). We first examined Otx2 and Visinin expression to confirm the reporter-observed effects, as these endogenous markers are expressed in both photoreceptors and restricted RPCs (Buenaventura et al., 2018; Emerson et al., 2013). In retinas with introduction of the CAG::MAML-DN construct, there was a significant increase of electroporated Visinin and Otx2-labeled cells, consistent with the reporter results (Fig. 3A, B). Olig2 expression, which is expected to be expressed in ThrbCRM1-labeled RPCs and not cones or HCs, was significantly increased, which suggests that the ThrbCRM1 reporter-positive population observed at this 2 day timepoint represents the restricted RPC state (Fig. 3A and 3C) (Emerson et al., 2013; Ghinia Tegla et al.; Hafler et al., 2012). In agreement with this observation and the ThrbCRM2 reporter result, expression of the cone gene Lhx4 was not qualitatively changed (Fig. S3B) (Buenaventura et al., 2019). In contrast, the representation of the H1 HC population (marked by Lim1 and AP2α; Fischer et al., 2007), normally generated by ThrbCRM1 restricted RPCs, is decreased in response to CAG::MAML-DN (Figure 3B). No significant changes in the H2-4 HC/RGC marker Isl1 or the Brn3 family of RGC markers were observed (Fig. S4) (Fischer et al., 2007; Liu et al., 2000). Taken together, these data suggest that the primary effect of Notch inhibition at this timepoint is to increase the formation of the restricted RPCs that generate cones and HCs.

**Fig. 3.**
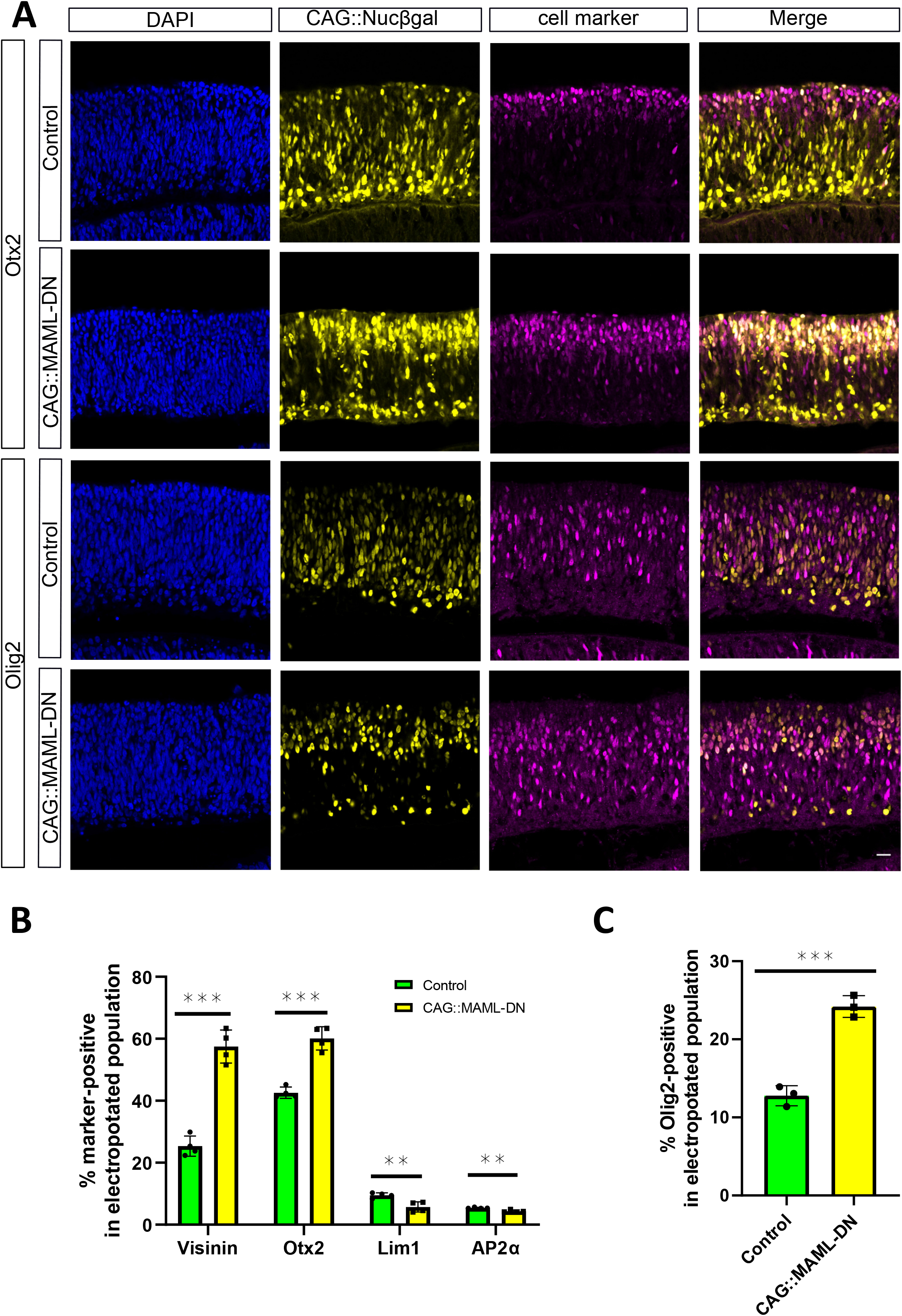
Effects of CAG::MAML-DN on endogenous retinal markers **A**. Confocal images of vertically sectioned E5 chick retinas co-electroporated with CAG::Nucßgal, and with or without CAG::MAML-DN and then cultured for two days. Sections were immunostained with Otx2 or Olig2 markers (magenta), CAG::Nucßgal (yellow), and nuclei visualized with DAPI. The scale bar shown in the bottom right picture denotes 40 µm and applies to all images. All images are oriented with the scleral side of the retina at the top of the image. **B**. Flow cytometry quantification of the percentage of Visinin, Otx2, Lim1 and AP2α-positive cells within the electroporated cells. Dissociated chick retinal cells electroporated *ex vivo* at E5 with the co-electroporation control CAG::TdTomato, under control (green bars) or CAG::MAML-DN (yellow bars) conditions. The retinas were cultured for two days. The Shapiro-Wilk normality test was used to confirm the normal distribution. ** signifies p<0.01, *** signifies p<0.001 with a two-tailed student’s t-test. Each point represents one biological replicate. **C**. Quantification of the percentage of Olig2-positive cells within the electroporated cell population from cell counting. Sectioned chick retinas electroporated *ex vivo* at E5 with the co-electroporation control CAG::Nucßgal. The retinas were cultured for two days. The Shapiro-Wilk normality test was used to confirm the normal distribution. *** signifies p<0.001 with a two-tailed student’s t-test. Each point represents one biological replicate.

To further validate that these MAML-DN effects were related to Notch signaling, we used flow cytometry to assess the effects of dominant-negative forms of Rbpj on these same DNA reporters and cell markers. Similar effects were observed upon introduction of both Rbpj-EnR and R218H constructs which support the conclusion that they are a result of reduced Notch signaling (Fig. S5). However, it was observed that the reduction of HCs in response to the Rbpj-EnR constructs was more severe than with MAML-DN, which is consistent with the Notch-independent role for Rbpj to function in a complex with Ptf1a, a known regulator of HC genesis (Beres et al., 2006; Hori et al., 2008; Lelièvre et al., 2011).

### Notch inhibition does not affect the number of cones produced from restricted RPCs

To test whether MAML-DN also induces a shift from HCs to cones within ThrbCRM1 reporter-positive RPCs, we replaced the ubiquitous CAG promoter upstream of MAML-DN with the ThrbCRM1 specific enhancer, so that MAML-DN would only be expressed in ThrbCRM1-active restricted RPCs. In contrast to CAG::MAML-DN, introduction of this ThrbCRM1::MAML-DN into E5 retinas did not alter the number of ThrbCRM1::GFP labeled RPCs. This suggests that the ectopic ThrbCRM1 cells observed with the CAG promoter are not due to induction of increased proliferation within the ThrbCRM1 population. Instead, this supports a role for Notch in the multipotent RPC population, in which the ThrbCRM1 element is not active (Fig. 4A).

**Fig. 4.**
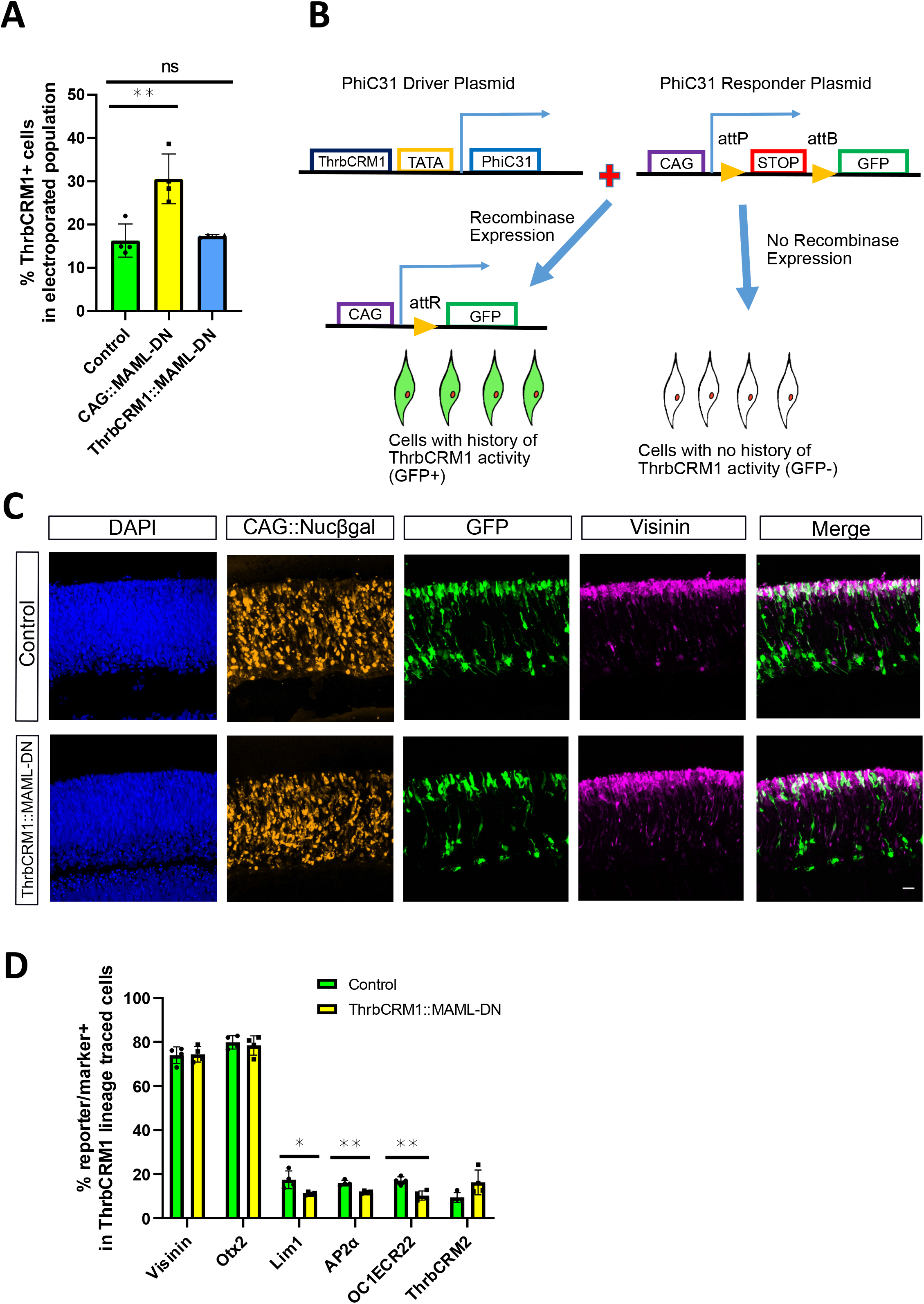
Targeted expression of MAML-DN to cone/HC restricted RPCs does not alter cone formation **A**. Flow cytometry quantification of the percentage of ThrbCRM1::GFP reporter-positive cells within all the electroporated cells. Dissociated chick retinal cells electroporated *ex vivo* at E5 with the CAG::iRFP co-electroporation control, and with or without CAG::MAML-DN or ThrbCRM1::MAML-DN and cultured for two days. The Shapiro-Wilk normality test was used to confirm the normal distribution. ANOVA with a post hoc Dunn test was used to test significance. ** signifies p<0.01, ns signifies p>0.05. **B**. Schematic representation of PhiC31 recombinase-based lineage tracing strategy (Schick et al., 2019). **C**. Confocal images of vertically sectioned E5 chick retinas co-electroporated with CAG::Nucßgal using ThrbCRM1::MAML-DN lineage trace with a PhiC31 responder plasmid shown in A. The retinas were cultured for two days post-electroporation. Sections were immunostained with Visinin (magenta), CAG::Nucßgal (orange), and nuclei visualized with DAPI. The scale bar shown in the bottom right picture denotes 40 µm and applies to all images. All images are oriented with the scleral side of the retina at the top of the image. **D**. Flow cytometry quantification of the percentage of Otx2, Lim1, AP2α, OC1ECR22 and ThrbCRM2 reporter-positive cells within the ThrbCRM1 lineage traced cell population. Dissociated chick retinal cells electroporated *ex vivo* at E5 with the co-electroporation control CAG::TdTomato, ThrbCRM1::PhiC31 and its responder plasmid, and with or without CAG::MAML-DN. The retinas were cultured for two days post-electroporation. The Shapiro-Wilk normality test was used to confirm the normal distribution. Mann-Whitney test was used to test significance in ThrbCRM2 quantification. A two-tailed student’s t-test was used to test significance in the rest of the quantifications. * signifies p<0.05, ** signifies p<0.01. Each point represents one biological replicate.

To genetically maintain labeling in post-mitotic daughter cells, we used a lineage tracing strategy to identify the retinal cells that have a ThrbCRM1 history (Fig. 4B) which were previously shown to be predominantly cones and H1 HCs (Schick et al., 2019). A driver plasmid was used in which the PhiC31 recombinase is transcriptionally regulated by the cis-regulatory element ThrbCRM1. In the responder plasmid, there is a STOP codon between a broadly active CAG promoter and GFP, which is flanked by two recombinase target sites. Excision of the STOP sequence upon ThrbCRM1-driven expression of PhiC31 recombinase, will lead to GFP expression in cells with a history of ThrbCRM1 activity (Schick et al., 2019).

Confocal analysis of retinas electroporated with the lineage tracing plasmids did not show any noticeable changes in the distribution of the electroporated cells or the amount of Visinin (Fig. 4C). Flow cytometry was used to quantify the percentage of cones and HCs within the GFP-labeled retinal cells after two days culture. Intriguingly, the number of cone photoreceptors did not change, as assessed by the expression of Visinin and Otx2, while the percentage of Lim1 and AP2α-labeled HCs was reduced (Fig. 4D and Fig. S6A). A small percentage of Isl1-labeled HCs and Brn3-labeled RGCs are formed from ThrbCRM1 restricted RPCs (Schick et al., 2019), and no significant changes in the populations were detected through cell counting in response to introduction of the ThrbCRM1::MAML-DN (Fig. S6B). Thus, the observed reduction in HCs did not correspond to increased formation of cone photoreceptors or Isl1-labeled HCs and RGCs in the daughter cell population of ThrbCRM1 restricted RPCs. These unidentified cells may be other cell types that cannot be identified by the cell markers used.

## Effects of MAML-DN on retinal cell proliferation and multipotency

Previous analysis of mouse Notch1 conditional knockout retinas identified a reduction in BrdU-positive cells, which suggested that loss of Notch signaling promoted premature cell cycle exit and down-regulation of cell proliferation (Jadhav et al., 2006b; Yaron et al., 2006). However, the difference in the percentages of BrdU-positive cells in wild-type and Notch1 CKO mutant is relatively small and the reason for this effect is not known. To further investigate the role of Notch in retinal cell proliferation, we co-electroporated CAG::MAML-DN and CAG::TdT into E5 chick retinas and after two days culture pulsed with EdU for 1 hour to label proliferating cells. We first quantified the percentage of EdU/CAG::TdT double-positive cells in all the electroporated cells using flow cytometry. Intriguingly, the population of proliferating cells does not significantly change compared to the control (Fig. 5A). Introduction of the Rbpj dominant-negative construct resulted in similar effects on electroporated EdU-positive cells (Fig. S7).

**Fig. 5.**
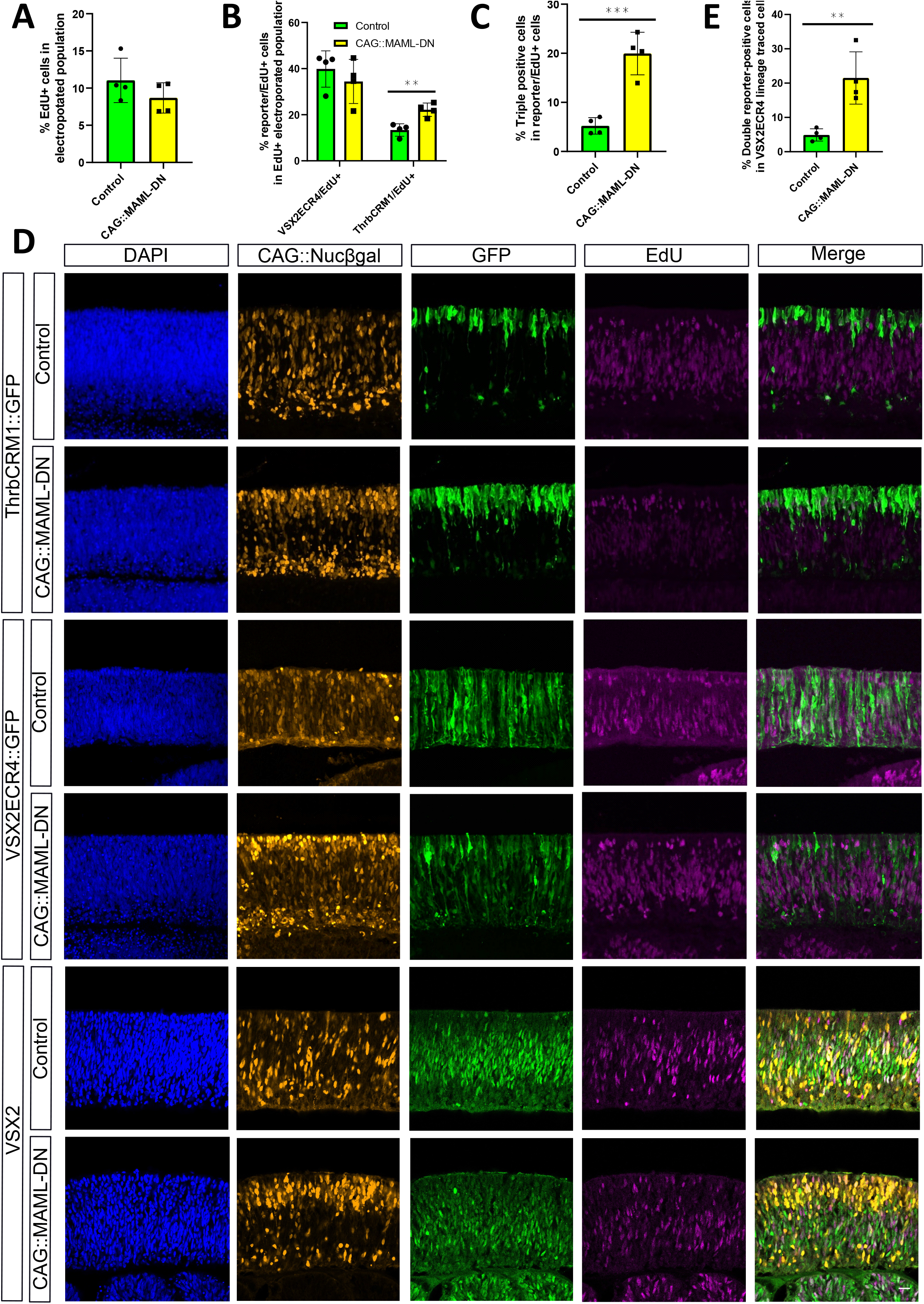
CAG::MAML-DN induced Notch inhibition leads to a shift from multipotent to cone/HC restricted RPCs **A**. Flow cytometry quantification of the percentage of EdU-positive cells within the electroporated cell population. Dissociated chick retinal cells electroporated *ex vivo* at E5 with the co-electroporation control CAG::TdTomato with or without CAG::MAML-DN. The retinas were cultured for two days after electroporation and pulsed with EdU for 1 hour before harvest. The Shapiro-Wilk normality test was used to confirm the normal distribution. A two-tailed student’s t-test was used to test significance. **B**. Flow cytometry quantification of EdU+ cells positive for VSX2ECR4::GFP or ThrbCRM1::TdTomato reporters. Dissociated chick retinal cells electroporated *ex vivo* at E5 with VSX2ECR4::GFP and ThrbCRM1::TdTomato. The retinas were cultured for two days after electroporation and pulsed with EdU for 1 hour before harvest. The Shapiro-Wilk normality test was used to confirm the normal distribution. Mann-Whitney test was used to test significance in VSX2ECR4::GFP/EdU+ quantification. A two-tailed student’s t-test was used to test significance in ThrbCRM1::TdTomato/EdU+ quantification. ** signifies p<0.01. **C**. Flow cytometry quantification of ThrbCRM1::TdTomato reporter-positive cells that also express VSX2ECR4::GFP reporter and EdU from dissociated chick retinal cells electroporated *ex vivo* at E5 with VSX2ECR4::GFP and ThrbCRM1::TdTomato. Retinas were cultured for two days and pulsed with EdU for 1 hour prior to harvest. The Shapiro-Wilk normality test was used to confirm the normal distribution. *** signifies p<0.001 with a two-tailed student’s t-test. **D**. Confocal images of vertically sectioned E5 chick retinas co-electroporated with CAG::Nucßgal, ThrbCRM1::GFP or VSX2ECR4::GFP, and with or without CAG::MAML-DN. The retinas were cultured for two days after electroporation and pulsed with EdU for 1 hour before harvest. The sections were immunostained with VSX2 maker (green), CAG::Nucßgal (orange), and nuclei visualized with DAPI. The scale bar shown in the bottom right picture denotes 40 µm and applies to all images. All images are oriented with the scleral side of the retina at the top of the image. **E**. Flow cytometry quantification of the percentage of VSX2ECR4/ThrbCRM1 double reporter-positive cells within the VSX2ECR4 lineage traced population. Dissociated chick retinal cells electroporated *ex vivo* at E5 with the co-electroporation control CAG::iRFP, VSX2ECR4::PhiC31 and its responder plasmid, and with or without CAG::MAML-DN. The retinas were cultured for two days post-electroporation. The Shapiro-Wilk normality test was used to confirm the normal distribution. ** signifies p<0.01 with a two-tailed student’s t-test. Each point represents one biological replicate.

These data suggest that Notch signaling does not directly control cell proliferation in retina. Although inhibition of Notch does not drastically alter the overall number of RPCs within this timeframe, the experiments described above suggest that it results in a shift from multipotent RPCs (decreased VSX2ECR4-positive cells) to restricted RPCs (ThrbCRM1-positive cells) with a significant increase in the number of ThrbCRM1/VSX2ECR4 double reporter-positive cells (Fig. 2B). To confirm that these reporter-positive populations are indeed proliferating, we co-electroporated ThrbCRM1 and VSX2ECR4 reporters and pulsed with EdU just prior to harvest to label RPCs. Flow cytometry was used to quantify the EdU/ThrbCRM1 and EdU/VSX2ECR4 double-positive cells in the electroporated population. We also measured the percentage of EdU/ThrbCRM1/VSX2ECR4 triple-positive cells versus the combination of EdU/ThrbCRM1 and EdU/VSX2ECR4 double-positive cells. There was a significant up-regulation of ThrbCRM1+/EdU+ cells as well as ThrbCRM1+/VSX2ECR4+/EdU+ cells that indicated a transition from VSX2ECR4-active RPCs to ThrbCRM1-active RPCs within the EdU+ population (Fig. 5B and 5C). In addition, immunostaining with the multipotent RPC marker VSX2 showed a down-regulation of VSX2+/EdU+ cells and an up-regulation of ThrbCRM1+/EdU+ cells, validating a shift between VSX2+ RPCs and ThrbCRM1-active RPCs (Fig. 5D).

To further determine if the induced ThrbCRM1 RPCs were derived from multipotent RPCs, we used the VSX2ECR4 element that is active in this population to drive expression of PhiC31 recombinase and conduct lineage tracing of this population. The number of VSX2ECR4/ThrbCRM1 double reporter-positive cells increased 5-fold in response to CAG::MAML-DN (Fig. 5E). This increase was consistent with the majority of new ThrbCRM1 reporter-positive cells being derived from multipotent RPCs, which suggests that Notch inhibition specifically affects the transition from the multipotent to restricted RPC state.

### Temporal parameters of the transition from multipotent RPCs to restricted RPCs

We were interested in determining the timeframe with which multipotent RPCs transition to restricted RPCs. To test if the changes in multipotent RPC numbers could be detected during their first cell cycle after electroporation, we electroporated E5 chick retinas with VSX2ECR4::GFP, ThrbCRM1::TdT, the co-electroporation reporter CAG::iRFP, and either CAG::MAML-DN or control conditions. Retinas were harvested at 8 hours after electroporation, which is a time before multipotent RPCs would be expected to complete one cell cycle. There were no significant differences in the number of VSX2ECR4-marked multipotent RPCs or ThrbCRM1-marked restricted RPCs (Fig. 6A). In addition, the VSX2ECR4/ThrbCRM1 double reporter-positive cells were present in a very low level as observed in wild type in both MAML-DN and Rbpj-EnR mutants (Fig. 6B and Fig. S7). After 20 hours culture, the ThrbCRM1-active population started to increase slightly but not nearly as much as observed after two days (Fig. 6C). However, the transition from multipotent RPCs to restricted RPCs showed a small but significant up-regulation, as the VSX2ECR4/ThrbCRM1 double reporter-positive cells increased (Fig. 6C). Examination of Otx2 and VSX2 endogenous markers revealed a more robust response, with a decreased VSX2-positive population and an increased Otx2-positive one after 20 hours culture (Fig. 6D). Taken together, these data suggest that the RPC transition induced by Notch inhibition did not occur at 8 hours post-electroporation, but could be detected after 20 hours, and increased further at 2 days post-electroporation. This suggests that multipotent RPCs do not immediately transition to a restricted RPC state, but do so after a delay that could be accounted for by a required progression through a single, or multiple rounds, of cell division.

**Fig. 6.**
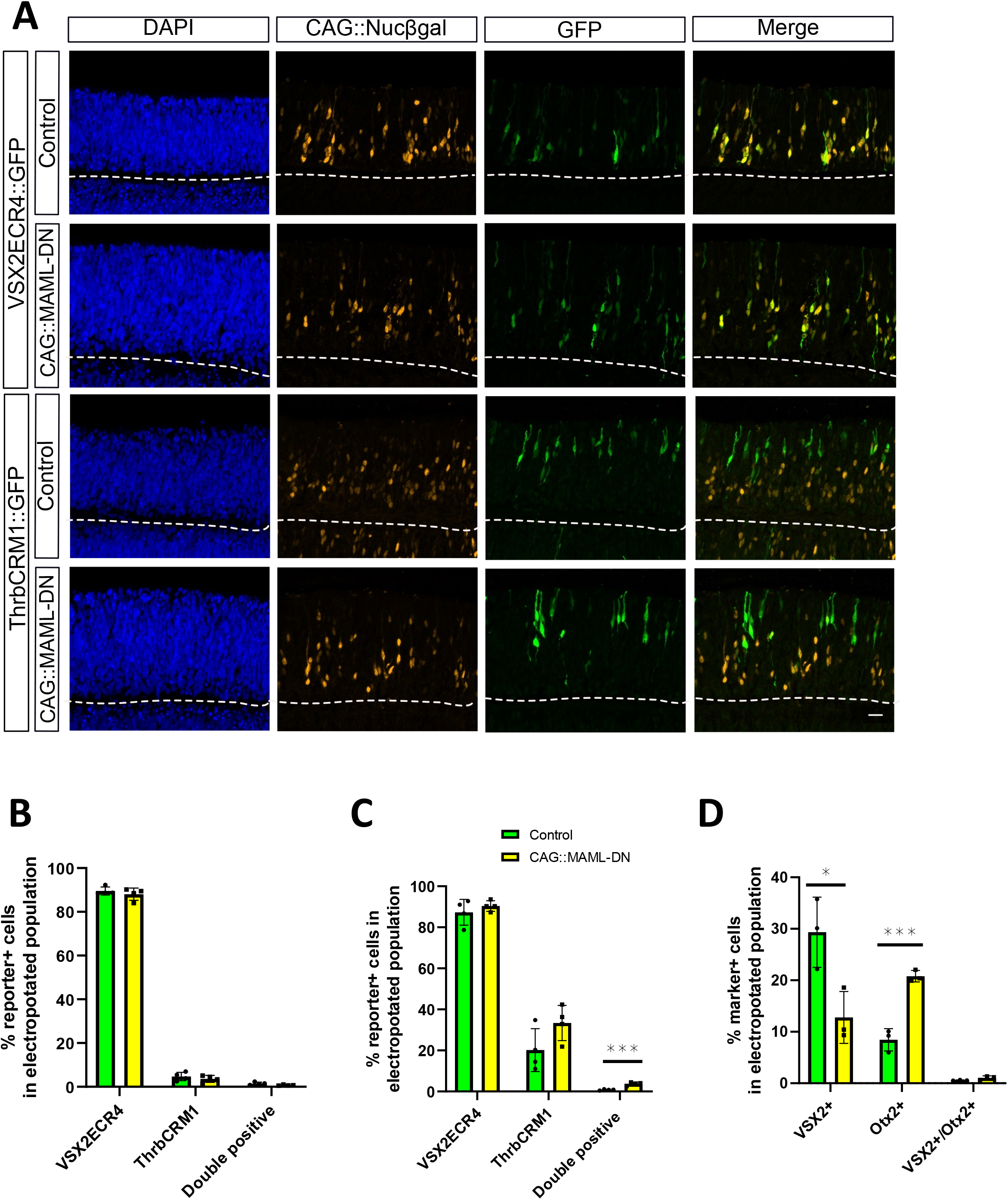
The short-term effects of Notch inhibition on RPC reporters **A**. Confocal images of vertically sectioned E5 chick retinas co-electroporated with CAG::Nucßgal, ThrbCRM1::GFP or VSX2ECR4::GFP, and with or without CAG::MAML-DN. The retinas were cultured for 8 hours post-electroporation. The scale bar shown in the bottom right picture denotes 40 µm and applies to all images. All images are oriented with the scleral side of the retina at the top of the image. **B**. Flow cytometry quantification of the percentage of VSX2ECR4, ThrbCRM1 and VSX2ECR4/ThrbCRM1 reporter-positive cells within all the electroporated cells. Dissociated chick retinal cells electroporated *ex vivo* at E5 with the co-electroporation control CAG::iRFP with or without CAG::MAML-DN. The retinas were cultured for 8 hours post-electroporation. The Shapiro-Wilk normality test was used to confirm the normal distribution. Mann-Whitney test was used to test significance in VSX2ECR4 quantification. A two-tailed student’s t-test was used to test significance in ThrbCRM1 and double-positive quantifications. **C**. Flow cytometry quantification of the percentage of VSX2ECR4, ThrbCRM1 and VSX2ECR4/ThrbCRM1 reporter-positive cells within all the electroporated cells. Dissociated chick retinal cells electroporated with CAG::iRFP *ex vivo* at E5 and cultured for 20 hours. The Shapiro-Wilk normality test was used to confirm the normal distribution. *** signifies p<0.001 with a two-tailed student’s t-test. Each point represents one biological replicate. **D**. Quantification of the percentage of VSX2, Otx2 and double marker-positive cells within the electroporated cell population from cell counting. Sectioned chick retinas electroporated *ex vivo* at E5 with the co-electroporation control CAG::Nucßgal with or without CAG::MAML-DN. The retinas were cultured for 20 hours and immunostained with VSX2 or Otx2 antibodies. The Shapiro-Wilk normality test was used to confirm the normal distribution. * signifies p<0.05, *** signifies p<0.001 with a two-tailed student’s t-test. Each point represents one biological replicate.

### Increased cone genesis in response to Notch inhibition is delayed relative to restricted RPC formation

While multiple experiments support the conclusion that Notch inhibition leads to the formation of an increased restricted RPC population, we did not detect any increased cone genesis within a 2 day post-electroporation window (Fig. 2B and Fig. S3). To investigate the possibility that there is a delay in cone genesis that will occur from the induced cone/HC RPCs, we used a lineage tracing strategy to follow the fates of multipotent RPCs and assessed the cone fate within this population. Chick retinas were electroporated with a VSX2ECR4::PhiC31 lineage tracing plasmid, a GFP lineage tracing reporter, a ThrbCRM2::TdTomato reporter, and CAG::iRFP to mark all electroporated cells (Fig. 7A). Retinal samples were harvested after two to five days culture. The total number of ThrbCRM2 reporter-positive cones did not change at 2 or 3 days post-electroporation, however, the number of ThrbCRM2 reporter-positive cells within the VSX2ECR4 lineage-traced population showed a significant increase at 3 days, in line with the hypothesis that the effects of Notch inhibition originate from multipotent RPCs (Fig. 7B). Both the overall ThrbCRM2 population and the population specifically lineage-traced from multipotent RPCs showed a very significant enrichment at 4 and 5 days after electroporation (Fig. 7C), which correlated in a delayed manner with the changes of ThrbCRM1 restricted RPCs from 1 to 2 days (Fig. 6C and Fig. 2B). This supports our hypothesis that increased cone genesis has a delay of 2 days relative to the up-regulated ThrbCRM1 RPCs formed in response to Notch inhibition and that this increased cone cell number is ultimately caused by effects that originate in multipotent RPCs.

**Fig. 7.**
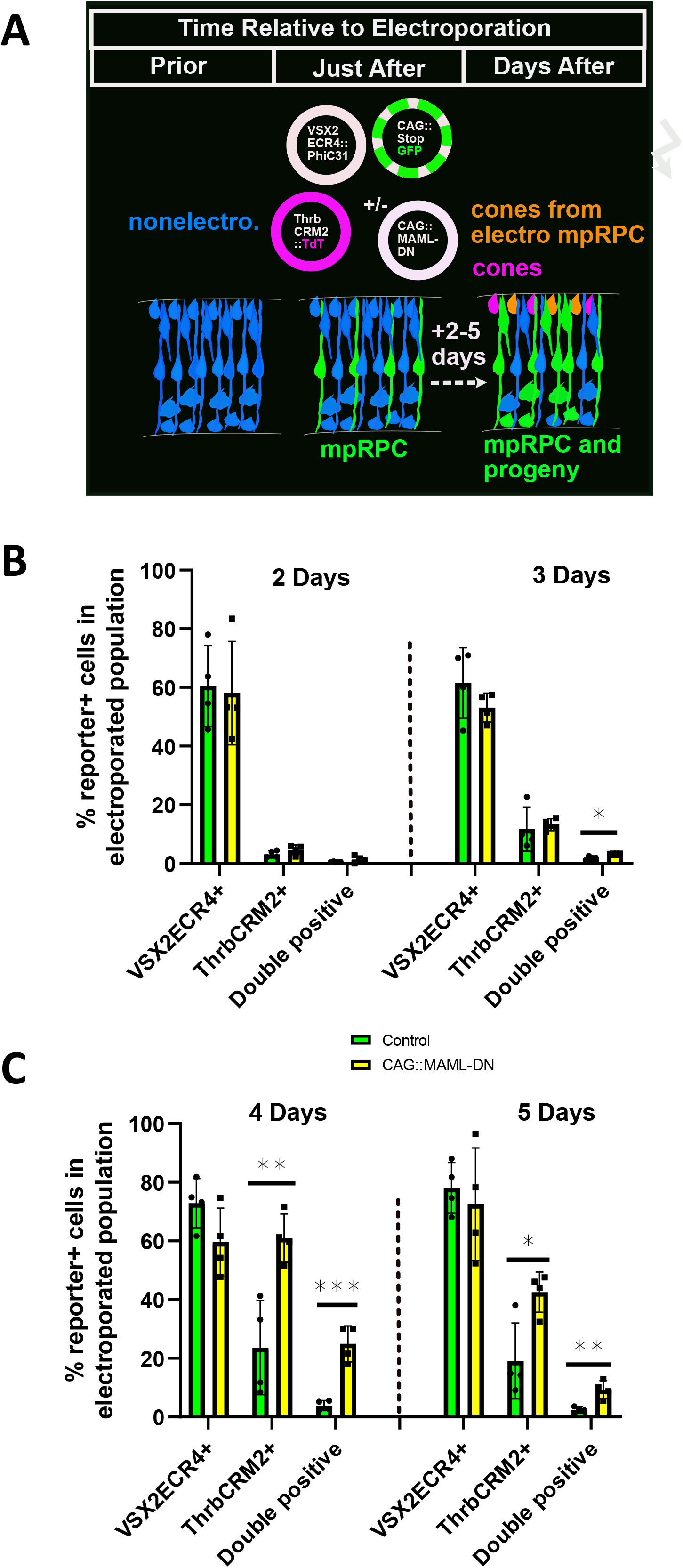
Notch inhibition leads to a delayed increase in cones relative to cone/HC RPCs **A**. Schematic of experimental set-up. Retinas were electroporated with VSX2ECR4::PhiC31 and CAG::stopGFP responder plasmid to label multipotent RPCs (mpRPC)s and their descendents. In addition, ThrbCRM2::TdT was co-electroporated to identify a subset of cones. CAG::MAML-DN was included in experimental conditions and omitted in control conditions. CAG::iRFP was included as a co-electroporation control (not shown). Retinas were assessed 2 to 5 days later by flow cytometry to measure GFP, TdT, and iRFP activity. Non-electroporated cells are shown in blue, lineage-traced cells other than cones in green, non-lineage traced ThrbCRM2+ cells in magenta and lineage traced ThrbCRM2+ cells in orange. **B**. Flow cytometry quantification of the percentage of ThrbCRM2::TdTomato reporter-positive cells within all the electroporated cells using VSX2ECR4 lineage trace strategy with the PhiC31 responder plasmid shown in Fig. 4B. Dissociated chick retinal cells electroporated *ex vivo* at E5 with the co-electroporation control CAG::iRFP, VSX2ECR4::PhiC31 and its responder plasmid, ThrbCRM2::TdTomato, and with or without CAG::MAML-DN. The retinas were cultured for 2 or 3 days post-electroporation. A two-tailed student’s t-test was used to test significance. The Shapiro-Wilk normality test was used to confirm the normal distribution. * signifies p<0.05, ** signifies p<0.01, *** signifies p<0.001 with a two-tailed student’s t-test. Each point represents one biological replicate. **C**. Same as A but with 4 or 5 days culture.

**Fig. 8.**
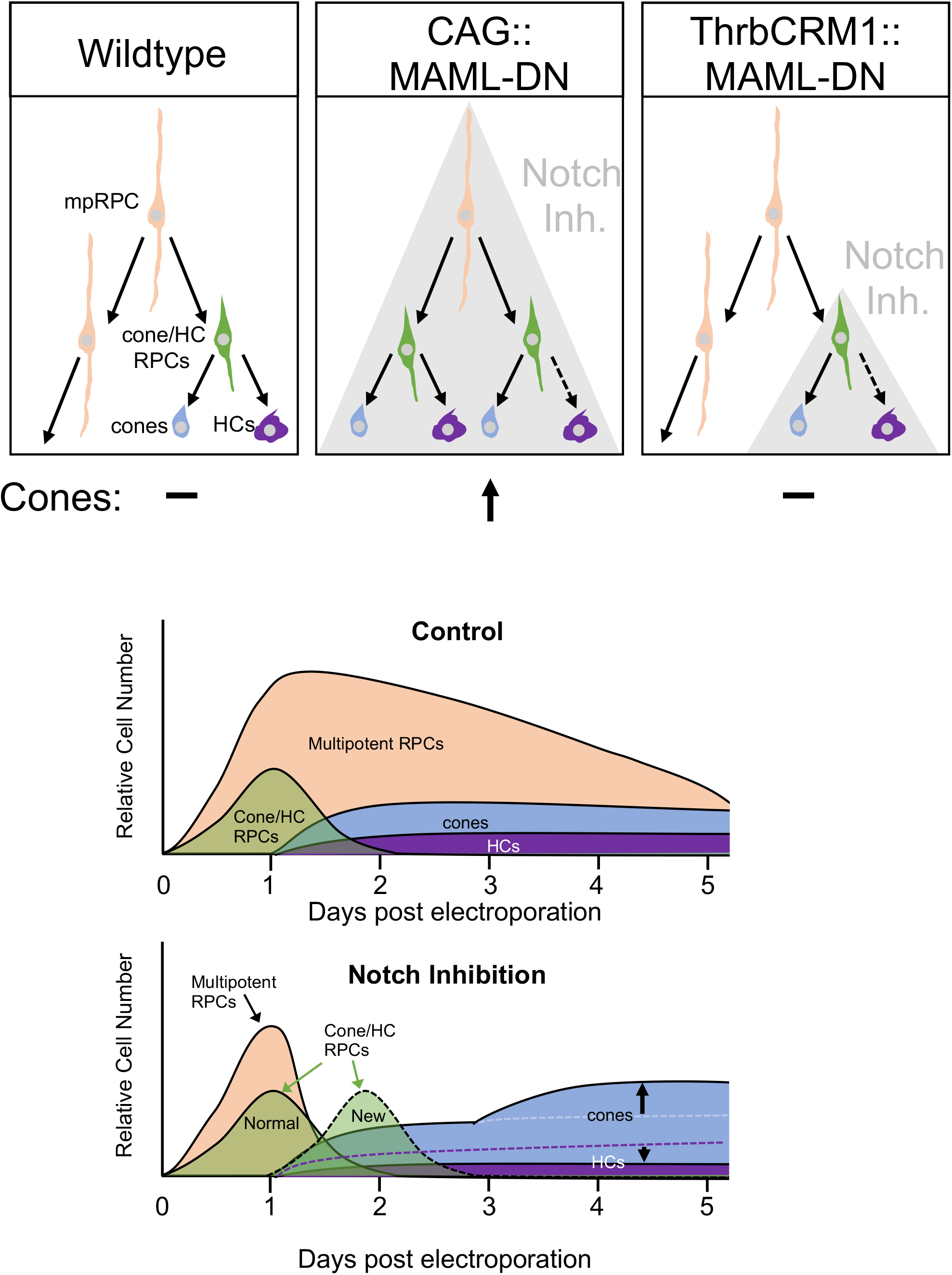
Model of Notch inhibition effects on cone/HC regulatory network Schematic of the general relationship between multipotent RPCs (mpRPC, beige), cone/HC RPCs (green), cones (blue), and HCs (purple). Grey shading denotes cell-type limits of Notch inhibition. CAG::MAML-DN is broadly active in all cells (denoted by grey shading) and leads to reduction in mpRPCs, increase in cone/HC RPCs and cones. Bottom graphs represent relative numbers of cell type populations under control and Notch inhibition conditions. Dotted lines represent decreased numbers.

## Discussion

While critical gene regulators of retinal cell type formation have been identified in the last decades, the relationships of these factors within gene regulatory networks and the cellular events in which they function are much less clear. One of the primary reasons for this is that, unlike other developmental systems, strict lineages of cell types originating from multipotent RPCs have not been identified, leading to some hypotheses that this system is driven primarily by stochastic choices (Gomes et al., 2011; He et al., 2012). However, identification of restricted RPC states with limited cell fate potential suggests that there are at least some deterministic processes involved and these are additional, discrete cellular steps in the formation of specific cell types during which genes could act. Here we attempted to integrate the well-documented role for the Notch pathway in the repression of cone photoreceptor formation and promotion of RPC proliferation, within a framework for retinal development that includes restricted RPCs.

We initially hypothesized that, similar to other systems, the effect of Notch inhibition on cone photoreceptors would be mediated through sister cell fate decisions. This would be consistent with the fact that there is a restricted RPC type that primarily forms cones and HCs and that several reports that examined the Notch1 mouse mutant identified an increase in cones and one also reported a decrease in HCs. However, though we also detected reduced HCs when we interfered with Notch signaling, we did not detect a concomitant increase in photoreceptors within the ThrbCRM1 restricted RPC daughter cell population. This suggests that sister cell fate choice is not the primary site of functional Notch signaling during cone photoreceptor formation. The most prominent and consistent effects of Notch signaling we observed were on the formation of the ThrbCRM1-active restricted RPC population and a concomitant depletion of the multipotent RPC population, which suggests that the primary role for Notch signaling in cone formation is at the step of restricted RPC derivation from multipotent RPCs. Our interpretation of these results is consistent with another recent developmental study that undertook a comprehensive transcriptome analysis of embryonic mouse retinas under conditions of Notch inhibition (Kaufman et al., 2019). Here, it was reported that the earliest effects of Notch inhibition through DAPT treatment include the loss of multipotent RPC gene expression (observed as well by earlier studies of Notch1 and Rbpj), but also the early increase of genes that have identified primary correlations with restricted RPCs and with reduced to no expression in postmitotic cones (Jadhav et al., 2006b; Riesenberg et al., 2009; Yaron et al., 2006). These gene expression changes include Olig2 (Hafler et al., 2012), Onecut1 (Emerson et al., 2013), and Otx2 (Emerson et al., 2013), all of which have been previously shown to be expressed in restricted RPCs that give rise to cones. Additional upregulated genes such as Dll4 (Rocha et al., 2009), and Atoh7 (Buenaventura et al., 2018), are expressed in cycling cells and are enriched in ThrbCRM1 cells as assessed by bulk RNA-Seq. All of these genes displayed a temporal dynamic consistent with increased production of restricted RPCs in this mouse model. In addition, several studies (Kaufman et al., 2019; Nelson et al., 2007) have identified an upregulation of several members of the bHLH family (NeuroD1, Ngn2, etc) which recent single cell analysis has associated with neurogenic RPCs and not multipotent RPCs (Lu et al., 2020).

A role for Notch signaling in maintaining a self-renewing multipotent RPC population is also consistent with its observed effects on proliferation. Many studies (Jadhav et al., 2006b; Kaufman et al., 2019; Yaron et al., 2006) have identified a decrease in cycling cells when Notch signaling is inhibited. We did not detect a significant decrease within two days of electroporation, but consistent with other analysis at early time points in the mouse, there was only a partial loss of cycling cells. Given the identified shift from multipotent to restricted RPCs identified here, we would predict that the majority of the remaining cycling cells in mouse Notch loss-of-function paradigms represent restricted RPCs and that the loss of multipotent RPCs is actually much larger than was previously assumed to be the case, as the existence of different RPC populations was not known at the time of those studies. The effects of Notch inhibition described here on RPC dynamics is also consistent with those observed during vertebrate forebrain development, where Notch inhibition leads to loss of apical progenitors and formation of intermediate progenitors, preceding the appearance of postmitotic neurons (Yoon et al., 2008).

The evidence presented in the current study, and interpreted from previous ones, suggests a primary role in Notch signaling during cone formation in mediation of the transition from multipotent to restricted RPC, but there are critical questions that remain to be addressed. One question is how cell division patterns of multipotent RPCs change with inhibition of Notch signaling? Instead of a multipotent RPC dividing asymmetrically to form another multipotent RPC and a ThrbCRM1 RPC, does Notch inhibition cause symmetrical divisions to form two ThrbCRM1 RPCs? A recent study in the zebrafish retina showed that pharmacological inhibition of Notch signaling led RPCs to transition from an asymmetric mode of division to a symmetric one in which two neurogenic RPCs were produced (Nerli et al., 2020). Our current findings align well with those of this study, with active Notch signaling functioning specifically to maintain multipotent RPCs and inhibit neurogenic (what we refer to as restricted) RPC states. Previous studies have also suggested that there may exist an additional RPC state located between multipotent RPCs and ThrbCRM1 RPCs (Schick et al., 2019; Suzuki et al., 2013). This RPC divides to form an RGC and a ThrbCRM1-like RPC on division. The formation of these cells could be the first change elicited by Notch inhibition which would suggest that RGCs should also increase and may correspond to the Ath5+ RPCs followed in Nerli et al., 2020. We do observe a number of cells in the RGC layer, however, we have not detected an increase in RGCs using early RGC markers such as Islet1 and Brn3. It is possible that the continued inhibition of the Notch signaling pathway in these cells interferes with their differentiation. This study focused on the formation of cone photoreceptors, but it will be interesting to determine if Notch functions at other temporal points in retinogenesis to control the formation of other restricted RPC types. Lastly, it will be informative to determine the direct functional targets of Notch signaling that drive RPC transitions. Identification of these pathways will have implications for our understanding of visual system evolution and for the development of strategies to control the proliferation dynamics and cell fate choices of stem cells for therapeutic uses.

## Methods

### Animals

All experimental procedures involving animals were carried out in consultation with the City College of New York, CUNY Institutional Animal Care protocols. Fertilized chick eggs were acquired from Charles River, stored in a 16 °C room for less than 10 days and incubated in a 38 °C humidified incubator for 5 days. All experiments that used animals were not sex-biased.

### DNA Plasmids

The Hes5::GFP reporter was provided by Dominique Henrique and described in (Vilas-Boas et al., 2011). CAG::Nucβgal was obtained from the Cepko lab (Cepko Lab, Harvard Medical School). Previously reported plasmids include CAG::TdT (Schick et al., 2019), OC1ECR22::GFP (Patoori et al., 2020; Schick et al., 2020), VSX2ECR4::GFP and CAG::iRFP (Buenaventura et al., 2018), ThrbCRM1::GFP and ThrbCRM2::GFP (Emerson et al., 2013), ThrbCRM1::PhiC31 and CaNa::GFP (Schick et al., 2019) and CAG::AU1Gapdh (Emerson and Cepko, 2011). pCAG is vector with a CAG promoter with no protein coding region downstream. To make the CAG::MAML-DN plasmid (Addgene plasmid # 160923, this study), an N-terminal MAML coding fragment (Weng et al., 2003) was amplified from chicken cDNA and fused with a mouse Gapdh coding sequence through overlap extension PCR. The PCR product was purified and digested with Age I /Nhe I before ligation into an Age I /Nhe I digested CAG::AU1Gapdh vector. ThrbCRM1::MAML-DN was made by replacing the CAG promoter with the ThrbCRM1 element. The CAG::Rbpj-EnR (Addgene plasmid # 160924, this study) and CAG::EnR-Rbpj (Addgene plasmid # 160925, this study) were made by fusing mouse Rbpj (GenBank ID BC051387.1 obtained from Transomic Technologies) with an engrailed repressor domain (Vickers and Sharrocks, 2002) and amplification through overlap extension PCR. The amplicon was digested by EcoR I and ligated into an EcoR I digested CAG vector. The R218H mutation was introduced into a mouse CAG::Rbpj plasmid through overlap extension PCR. All the plasmids described above were validated by restriction digest and Sanger sequencing.

### Electroporation and Culture

To deliver plasmid DNA into dissected retinas, *ex vivo* experiments were carried out as described in (Emerson and Cepko, 2011) using a Nepagene NEPA21 Type II Super Electroporator. DNA mixes for *ex vivo* electroporation were made in 1X PBS with a final concentration of 0.1 μg/μl for the co-electroporation control plasmids with CAG promoters and 0.16 μg/μl for reporter and dominant-negative plasmids. Retinas were dissected at E5 and cultured between 8 hours and 5 days after electroporation. In all experiments, “control” refers to samples in which the same concentration of pCAG plasmid was substituted for the experimental plasmid in the electroporation.

### Immunohistochemistry

After harvest, retinas were fixed in 4% paraformaldehyde for 30 minutes, washed 3 times in 1XPBS, and sunk in 30% sucrose/0.5XPBS overnight. Retinas were frozen in OCT (Sakura Tissue-Tek, 4583). 20 μm vertical sections were acquired using a Leica Cryostat and collected on 5-7 slides (FisherBrand, 12-550-15).

Sections were blocked in 5% serum (Jackson ImmunoResearch, Donkey - 017-000121, Goat - 005-000-121) in 1XPBT (0.01% Tween-20 in 1XPBS) for 1 hour at room temperature and incubated in primary antibody at 4°C overnight. The primary antibodies used were: chicken anti-GFP (ab13970, Abcam, 1:2000), rabbit anti-GFP (A-6455, Invitrogen, 1:1000), chicken anti-β-galactosidase (ab9361, Abcam, 1:1000), mouse IgG1 anti-β-galactosidase (40-1a-s, DSHB 1:20), rabbit anti-RFP (600-401-379, Rockland, 1:250), mouse IgG1 anti-Visinin (7G4-s, DSHB, 1: 500), mouse IgG1 anti-Lim1 (4F2-c, DSHB, 1:30), mouse anti-AP2α(3B5, DSHB, 1:200), mouse IgG1 anti-Islet1 (39.3F7, DSHB, 1:50), mouse IgG1 anti-Brn3a (MAB1585, EMD Millipore, 1:800), mouse anti-Pan Brn3 (Sc-390781, Santa Cruz, 1:100), sheep anti-Vsx2 (x1180p, Ex Alpha, 1:200), goat anti-Otx2 (AF1979, R&D, 1:500), rabbit anti-Otx2 (AB21990, Abcam, 1:500) and rabbit anti-Olig2 (4B9610-AF488, Millipore, 1:500). Sections were washed 3 times with 1XPBT, blocked at room temperature for 30 min and incubated in secondary antibody at 4°C overnight. The secondary antibodies used were from Jackson Immunoresearch and appropriate for multiple labeling. Alexa 488- and 647-conjugated secondary antibodies (resuspended in 50% glycerol) were used at 1:400 and cy3-conjugated antibodies were used at 1:250 as described (Schick et al., 2019). 4’6-diamidino-2-phenylindole (DAPI) was added at 1 μg/ml after 3X wash with 1XPBT. Sections were mounted using Fluoromount-G (Southern Biotech, 0100-01) and 34×60 mm coverslips (VWR, 48393-106).

### EdU Labeling

Before fixation, retinas were incubated at 37°C for 1 hour with 50 μM EdU in culture media and harvested as described above. Sections were permeabilized for 15 minutes in 0.1% Triton in 1X PBS before incubating with the EdU Reaction Cocktail for 30 minutes in the dark. Sections were washed 3 times with 1X PBT before blocking and antibody staining. EdU was detected using a Click-iT EdU Alexa Fluor 647 imaging kit (Invitrogen, C10340).

### Confocal Microscopy and Cell Counting

Confocal images were acquired using a Zeiss LSM710 inverted confocal microscope with an EC Plan-Neofluar 40x/1.30 Oil DIC M27 immersion objective, 488nm laser, 561 nm laser, 633 nm laser, 405 nm laser, and ZEN Black 2015 21 SP2 software resolution of 1024 x 1024, acquisition speed of 6. Fiji (Schindelin et al., 2012) was used to analyze images and convert into picture format. Cell counting was performed using the Cell Counter plugin-in for ImageJ developed by Kurt De Vos. Cell counts used at least 3 retinas for each condition, 3 sections per retina. Figures were assembled using Affinity Designer vector editor (Serif [Europe] Ltd). Brightness and contrast were adjusted uniformly in all the figures. All retinal sections are oriented with the developing photoreceptor side at the top of the image. All images are presented as maximum projections, with the exception of Olig2-related images in Figure 3A, which are single z-planes.

### Retina Dissociation and Flow Cytometry

After ex vivo culture, the remaining retinal pigment epithelium (RPE) and condensed vitreal materials were removed in HBSS (GIBCO, 14170112) and dissociated following a papain-based (Worthington, L5003126) protocol (Jean-Charles et al., 2018). Dissociated retinal cells were fixed using 4% paraformaldehyde for 15 minutes and washed 3 times in 1X PBS. Samples for quantification were filtered into 4ml tubes (BD Falcon, 352054) through 40um cell strainers (Biologix, 15-1040). Dissociated cells were analyzed with a BD LSR II flow cytometer using the 488nm, 561nm, and 633nm lasers. All flow cytometry data was analyzed using FlowJo Version 10.4.2.

### Statistical Analysis

Graphs and statistical tests were conducted using Microsoft Office Excel and GraphPad Prism 8. The error bars represent S.E.M. ANOVA with a post hoc Dunn test was used in Fig. 4A and Fig. S2B. Mann-Whitney test was used in Fig. 4D, Fig. 5B, Fig 6B, Fig. S4B and Fig. S7B. A two-tailed student’s t-test was used for all other statistical tests. All the tests were performed with independent samples after the Shapiro-Wilk normality test using GraphPad Prism8, and the results were significant as noted. Quantifications for flow cytometry were based on 4 biological replicates per condition and at least 2 technical replicates. Quantifications for cell counting were based on 3 biological replicates per condition with 3 sections per retina.

## Acknowledgements

Funding for this project was provided by NSF CAREER 1453044 (to M.E.) and supported by NIMHD 3G12MD007603-30S2 (to CCNY). Jeffrey Walker and Jorge Morales provided technical support with flow cytometry and confocal microcopy experiments. We thank Diego Buenaventura for providing the VSX2ECR4::PhiC31 plasmid and for guidance with flow cytometry analysis. We thank Miruna Ghinia-Tegla for advice on cell counting and Estie Schick for comments on the manuscript. Shirley Chan, Denice Moran and Boguslava Dimitrova provided technical assistance. We thank all the members of the Emerson lab for support throughout the project.

## Author Contributions

XC and ME designed the experiments and analyzed the data. XC and ME conducted the experiments. XC and ME wrote the manuscript.

## Figure Legends

**Fig. S1**. CAG::MAML-DN expression and validation in retinal cells

**A**. Confocal images of vertically sectioned E5 chick retinas co-electroporated with CAG::Nucßgal, Hes5::GFP, and with CAG::MAML-DN or empty AU1-Gapdh vector and cultured for two days before AU1 immunostaining. The scale bar shown in the bottom right picture denotes 40 µm and applies to all images. All images are oriented with the scleral side of the retina at the top of the image. **B**. Bar graphs of flow cytometry analysis of dissociated chick retinal cells electroporated *ex vivo* at E5 with the CAG::iRFP co-electroporation control, Hes5::GFP reporter, ThrbCRM1::TdT, and with CAG::MAML-DN or empty AU1-Gapdh vector and cultured for two days. The number of reporter-positive cells within the electroporated population is plotted. The Shapiro-Wilk normality test was used to confirm the normal distribution. *** signifies p<0.001 with a two-tailed student’s t-test. Each point represents one biological replicate.

**Fig. S2**. Rbpj-EnR and R218H have similar effects as CAG::MAML-DN on Hes5 Notch reporter

**A**. Confocal images of vertically sectioned E5 chick retinas co-electroporated with CAG::Nucßgal, Hes5::GFP Notch reporter, and with or without Rbpj-EnR and cultured for two days before immunostained with Visinin. The scale bar shown in the bottom right picture denotes 40 µm and applies to all images. All images are oriented with the scleral side of the retina at the top of the image.

**B**. Flow cytometry quantification of the percentage of Hes5::GFP-positive cells within all the electroporated cells. Dissociated chick retinal cells electroporated *ex vivo* at E5 with the CAG::iRFP co-electroporation control, Hes5::GFP, and with or without EnR-Rbpj or Rbpj-EnR and cultured for two days. The Shapiro-Wilk normality test was used to confirm the normal distribution. ANOVA with a post hoc Dunn test was used to test significance.*** signifies p<0.001. Each point represents one biological replicate. **C**. Same as A but with or without R218H.

**Fig. S3**. CAG::MAML-DN does not promote cone formation at two days post-electroporation

**A**. Confocal images of vertically sectioned E5 chick retinas co-electroporated with CAG::Nucßgal, ThrbCRM2::GFP or OC1ECR22::GFP, and with or without CAG::MAML-DN and cultured for two days. The scale bar shown in the bottom right picture denotes 40 µm and applies to all images. All images are oriented with the scleral side of the retina at the top of the image. **B**. Confocal images of vertically sectioned E5 chick retinas co-electroporated with CAG::Nucßgal, ThrbCRM1::GFP, and with or without CAG::MAML-DN and cultured for two days. LHX4 antibody was used to label cones. The scale bar shown in the bottom right picture denotes 40 µm and applies to all images. All images are oriented with the scleral side of the retina at the top of the image.

**Fig. S4**. CAG::MAML-DN does not shift the number of H2, H3 and H4 HCs and RGCs

**A**. Confocal images of vertically sectioned E5 chick retinas co-electroporated with CAG::Nucßgal, and with or without CAG::MAML-DN and cultured for two days. Sections were immunostained with Islet1 and Pan Brn3 cell specific markers (magenta), CAG::Nucßgal (orange), and nuclei visualized with DAPI. The scale bar in the bottom right panel denotes 40 µm and applies to all images. All images are oriented with the scleral side of the retina at the top of the image. **B**. Flow cytometry quantification of the percentage of Islet1, Brn3a and Pan Brn3-positive cells within all electroporated cells. Dissociated chick retinal cells electroporated *ex vivo* at E5 with the CAG::TdTomato co-electroporation control and cultured for two days. The Shapiro-Wilk normality test was used to confirm the normal distribution. A two-tailed student’s t-test was used to test significance in Islet1 and Brn3a quantifications. Mann-Whitney test was used to test significance in Pan Brn3 quantification. Each point represents one biological replicate.

**Fig. S5**. Rbpj-EnR and R218H have similar effects as CAG::MAML-DN on ThrbCRM1 RPCs

**A**. Flow cytometry quantification of the percentage of VSX2ECR4::GFP, ThrbCRM1::TdTomato, VSX2ECR4::GFP/ThrbCRM1::TdTomato double reporter-positive, Visinin, ThrbCRM2::TdTomato, OC1ECR22::GFP, Lim1 and AP2α-positive cells within all the electroporated cells. Dissociated chick retinal cells electroporated *ex vivo* at E5 with the CAG::iRFP or CAG::TdTomato co-electroporation control, and with or without Rbpj-EnR and cultured for two days. The Shapiro-Wilk normality test was used to confirm the normal distribution. ** signifies p<0.01, *** signifies p<0.001 with a two-tailed student’s t-test. **B**. Flow cytometry quantification of the percentage of ThrbCRM2::TdTomato, VSX2ECR4::GFP, ThrbCRM1::TdTomato, VSX2ECR4::GFP/ThrbCRM1::TdTomato double reporter-positive cells within all the electroporated cells. Dissociated chick retinal cells electroporated *ex vivo* at E5 with the CAG::iRFP co-electroporation control, and with or without R218H and cultured for two days. The Shapiro-Wilk normality test was used to confirm the normal distribution. ANOVA with a post hoc Dunn test was used to test significance. * signifies p<0.05. Each point represents one biological replicate.

**Fig. S6**. ThrbCRM1::MAML-DN does not have effects on H2, H3 and H4 HCs and RGCs within ThrbCRM1 lineage traced population

**A**. Confocal images of vertically sectioned E5 chick retinas co-electroporated with CAG::Nucßgal, ThrbCRM1::PhiC31 and its responder plasmid, and with or without ThrbCRM1::MAML-DN. The retinas were cultured for two days post-electroporation. The sections were immunostained with Otx2, Lim1 and AP2α markers (magenta), CAG::Nucßgal (orange), and nuclei visualized with DAPI. The scale bar shown in the bottom right picture denotes 40 µm and applies to all images. All images are oriented with the scleral side of the retina at the top of the image. **B**. Quantification of the percentage of Islet1 and Brn3 marker-positive cells within the ThrbCRM1 lineage traced cell population from cell counting. Sectioned chick retinas electroporated *ex vivo* at E5 with the co-electroporation control CAG::Nucßgal, ThrbCRM1::PhiC31 and its responder plasmid, and with or without ThrbCRM1::MAML-DN. The retinas were cultured for two days after electroporation. The Shapiro-Wilk normality test was used to confirm the normal distribution. A two-tailed student’s t-test was used to test significance. Each point represents one biological replicate.

**Fig. S7**. Rbpj-EnR induced Notch inhibition has similar effects as CAG::MAML-DN on cell proliferation

**A**. Flow cytometry quantification of the percentage of EdU-positive cells within all the electroporated cells. Dissociated chick retinal cells electroporated *ex vivo* at E5 with the co-electroporation control CAG::TdTomato with or without Rbpj-EnR. The retinas were cultured for two days after electroporation and pulsed with EdU for 1 hour before harvest. The Shapiro-Wilk normality test was used to confirm the normal distribution. A two-tailed student’s t-test was used to test significance. **B**. Flow cytometry quantification of the percentage of VSX2ECR4, ThrbCRM1 or VSX2ECR4/ThrbCRM1 reporter-positive cells within the electroporated cell population. Dissociated chick retinal cells electroporated *ex vivo* at E5 with the co-electroporation control CAG::iRFP, VSX2ECR4::GFP or ThrbCRM1::TdTomato, and with or without Rbpj-EnR. The retinas were cultured for 8 hours post-electroporation. The Shapiro-Wilk normality test was used to confirm the normal distribution. A two-tailed student’s t-test was used to test significance in ThrbCRM1 quantification. Mann-Whiteney test was used to test significance in VSX2ECR4 and double reporter-positive quantifications. * signifies p<0.05. Each point represents one biological replicate.

