## Supplementary figures and images for "Notch signaling represses cone photoreceptor formation through the regulation of retinal progenitor cell states"

### Supplemental Figure 1

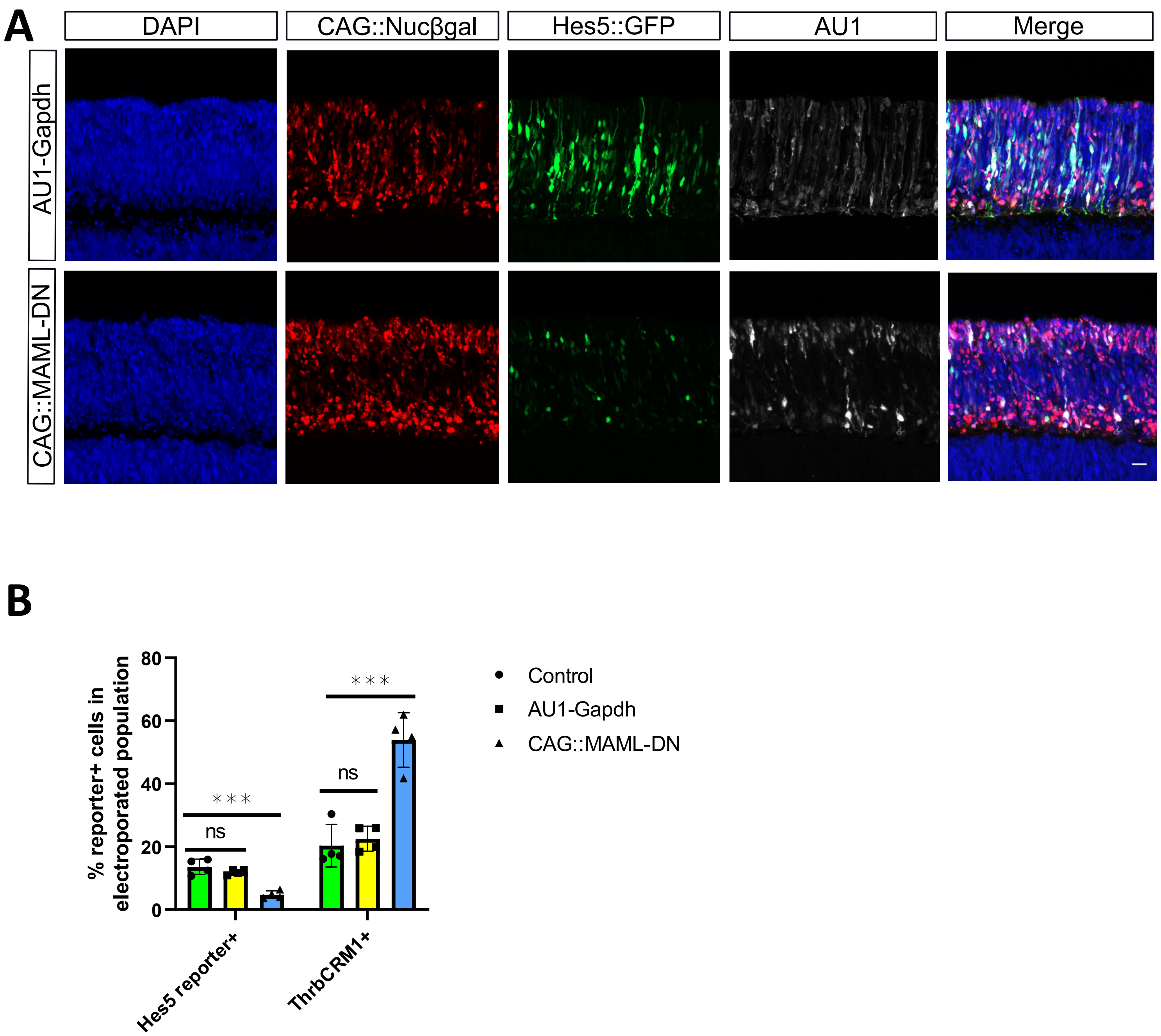

### Supplemental Figure 2

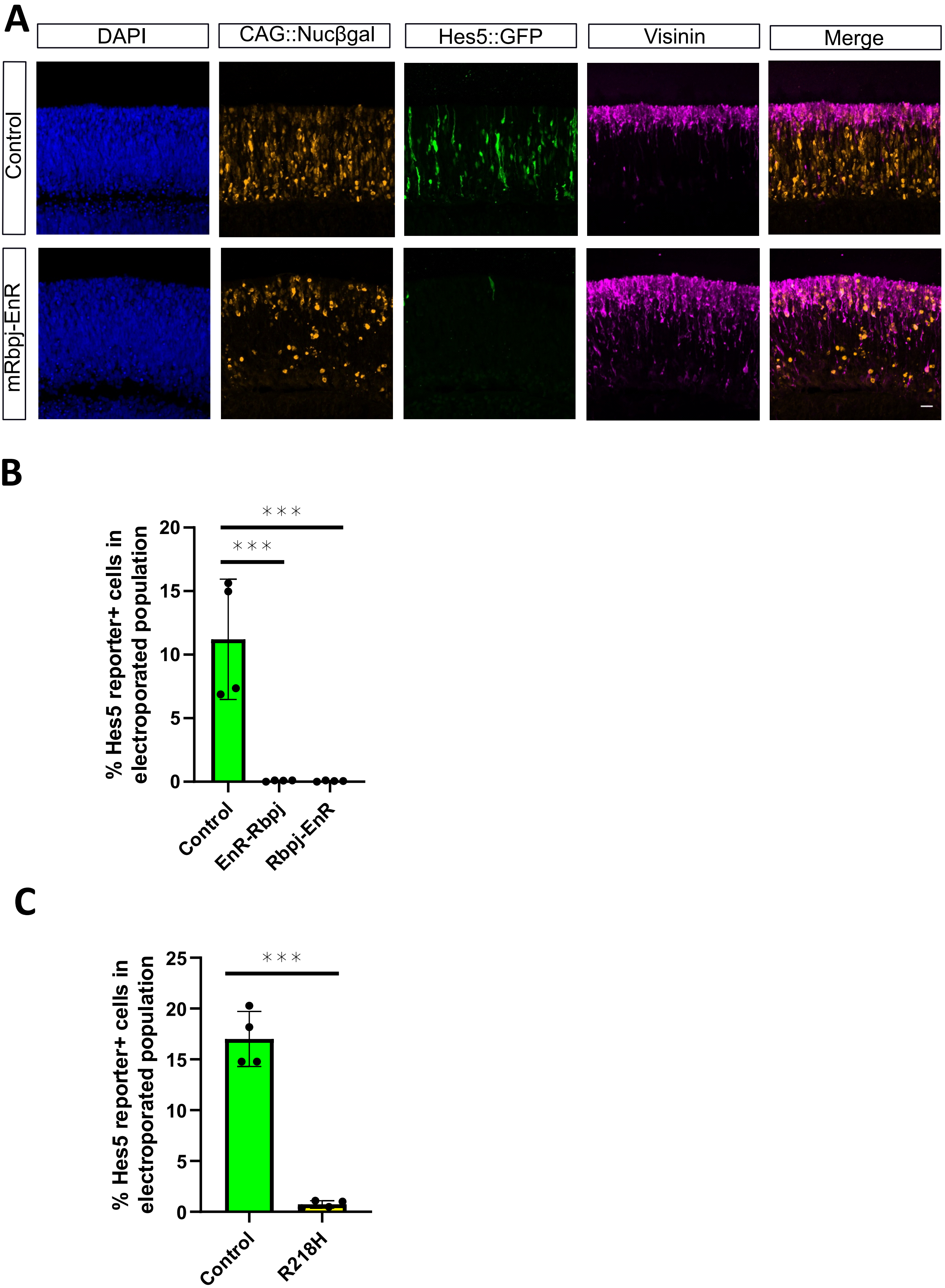

### Supplemental Figure 3

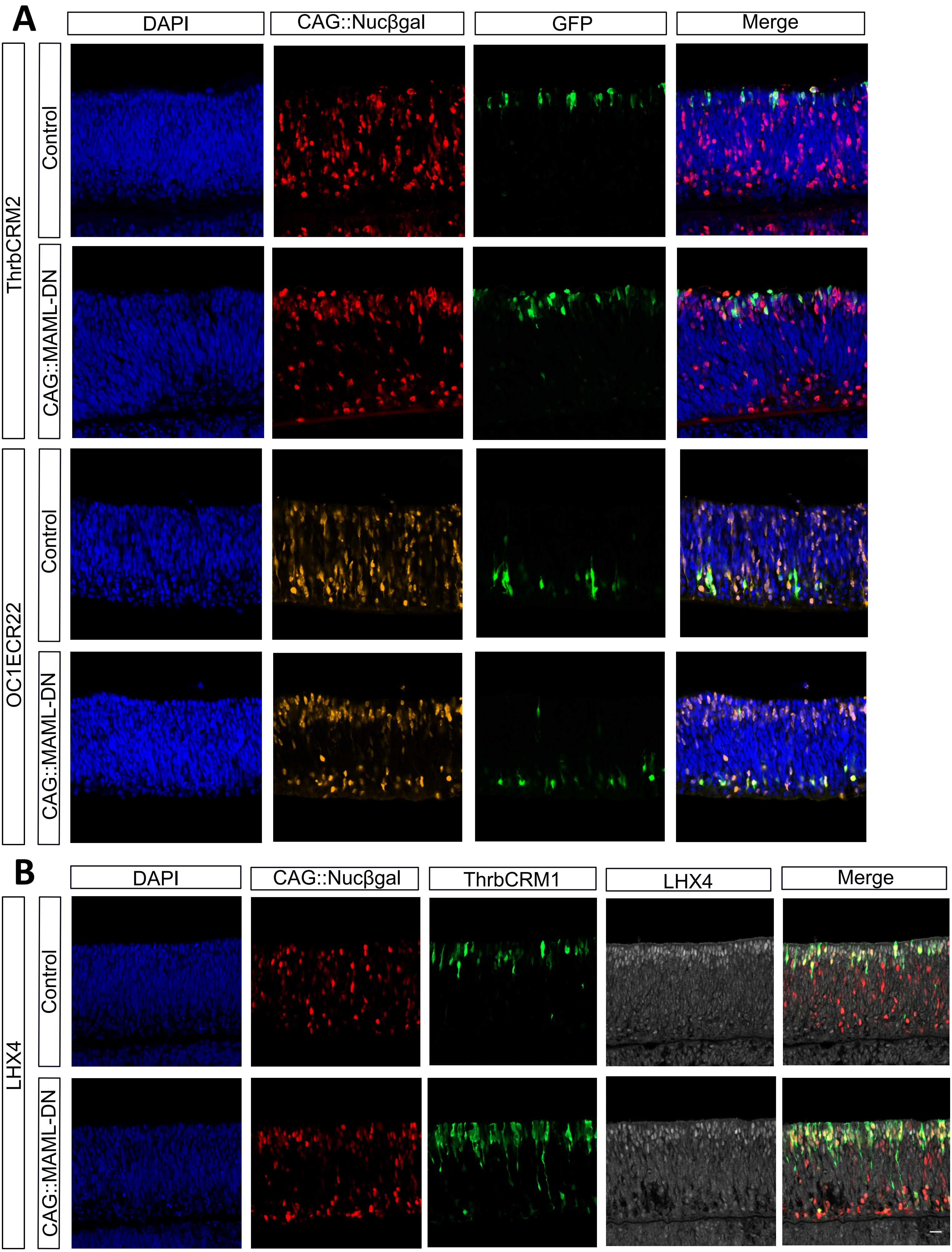

### Supplemental Figure 4

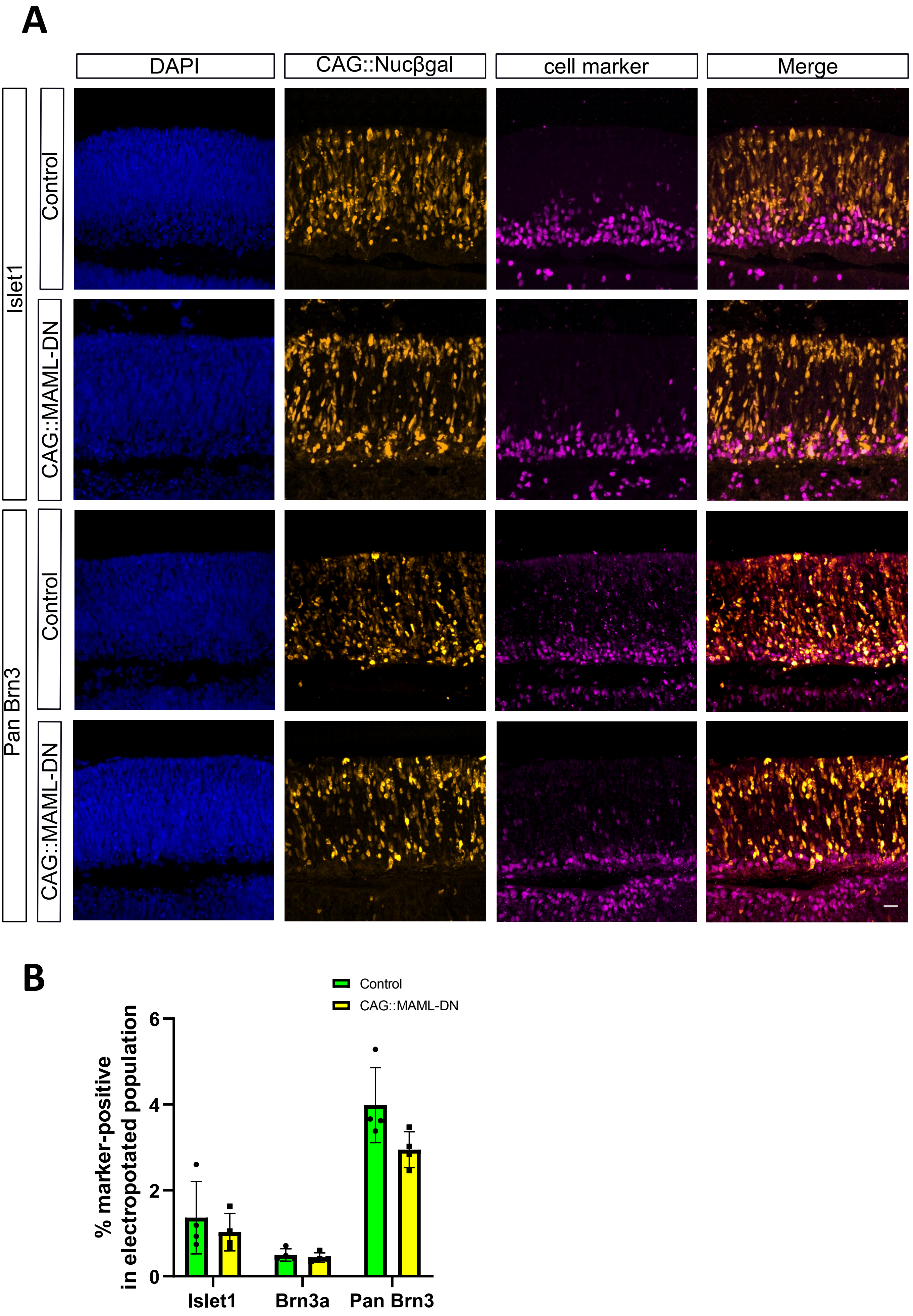

### Supplemental Figure 5

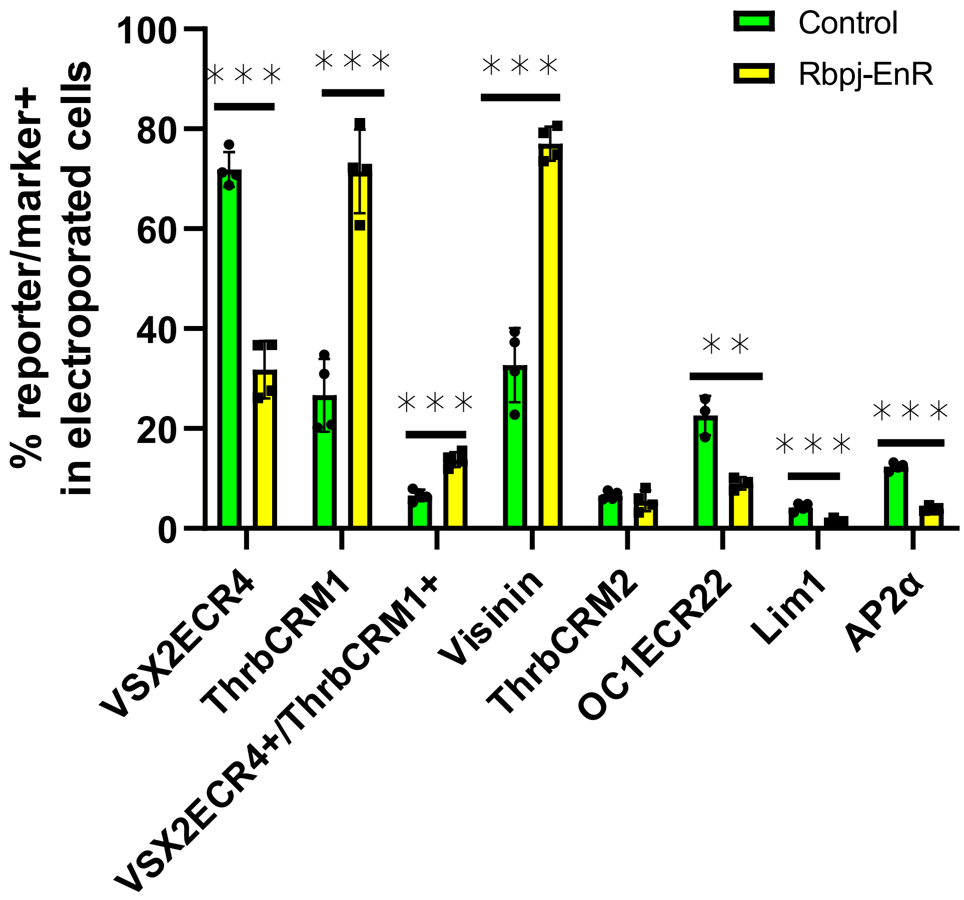

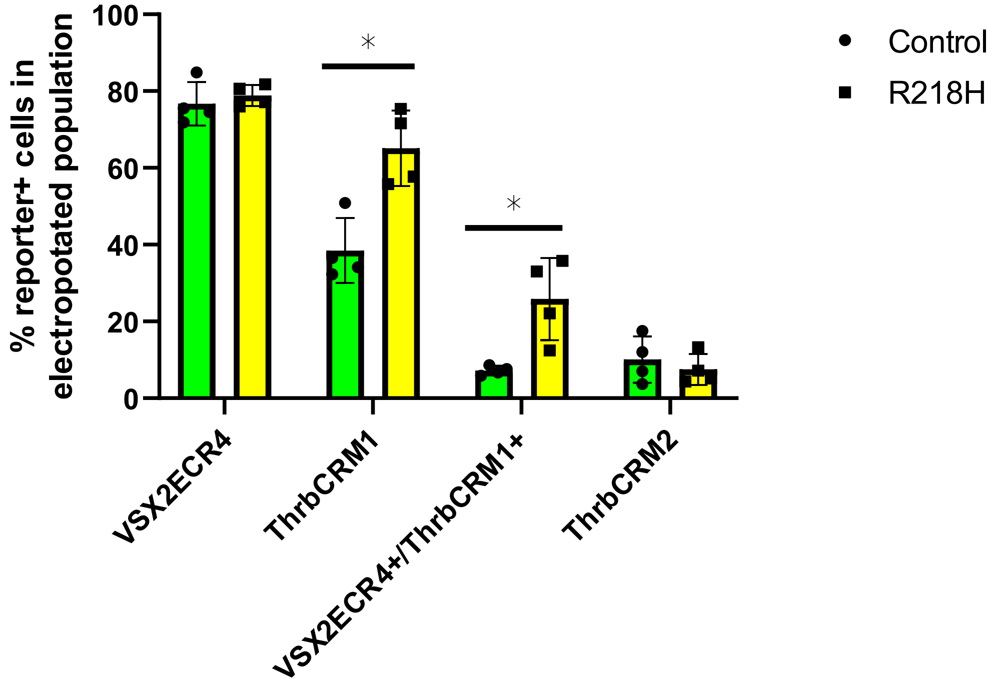

### Supplemental Figure 6

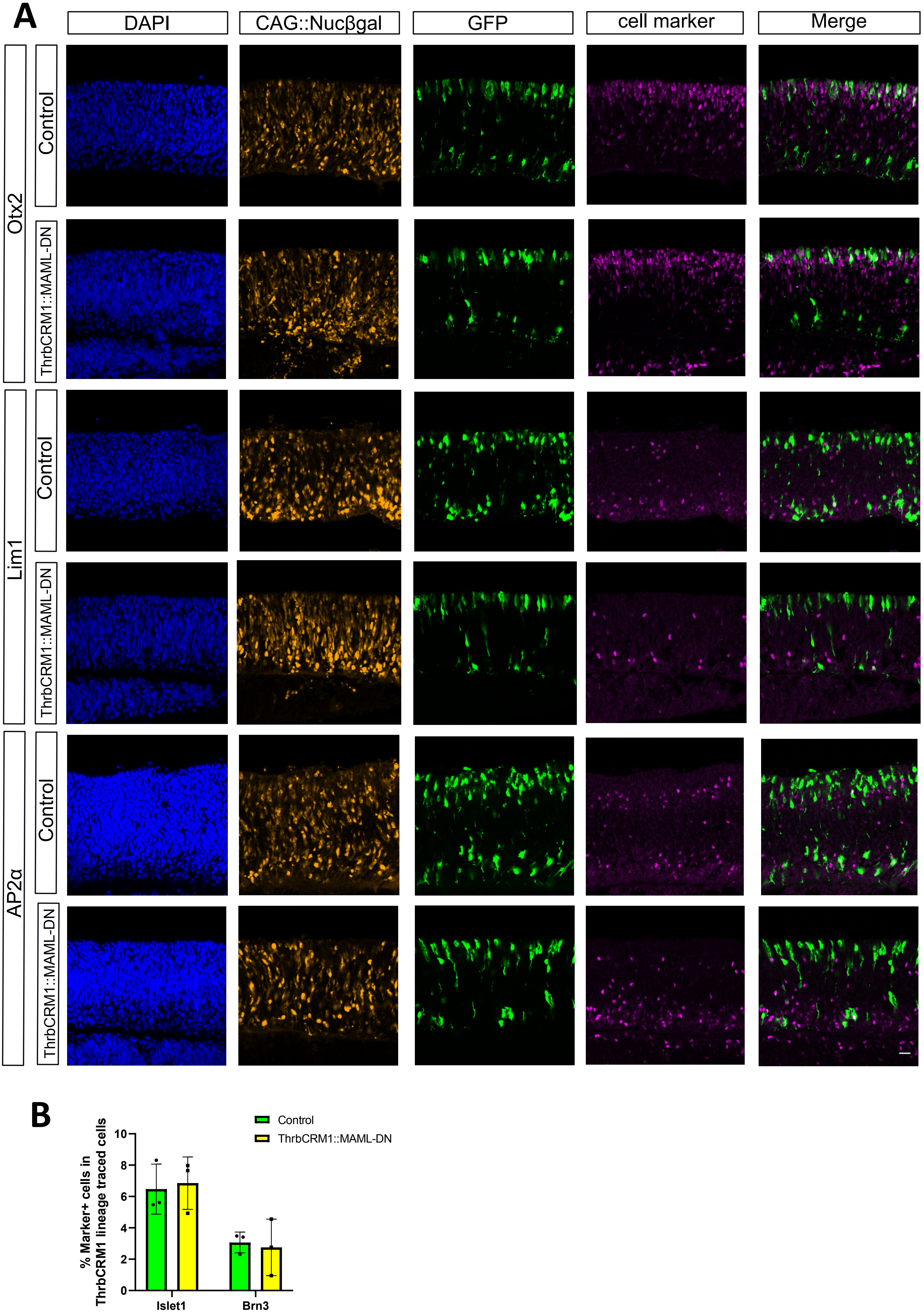

### Supplemental Figure 7

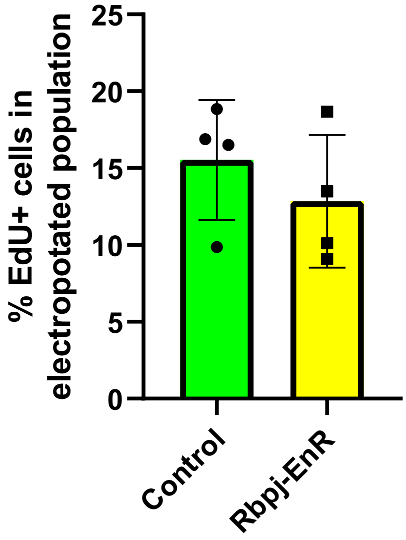

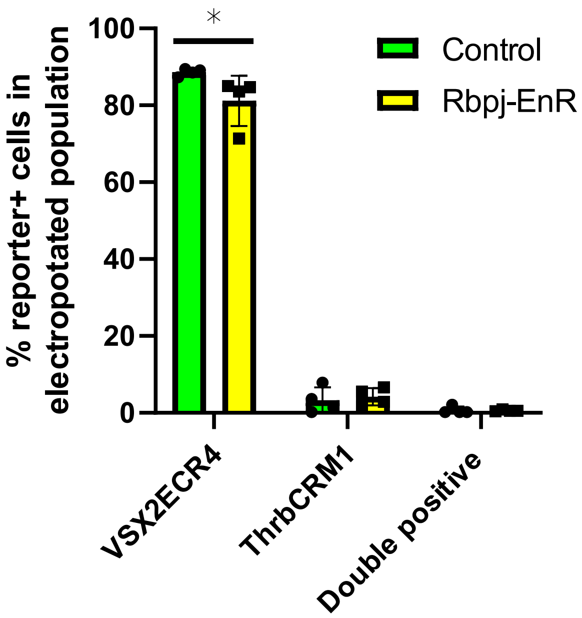
